# High throughput intracellular delivery by viscoelastic mechanoporation

**DOI:** 10.1101/2023.04.24.538131

**Authors:** Derin Sevenler, Mehmet Toner

## Abstract

Brief and intense electric fields (electroporation) and/or tensile stresses (mechanoporation) have been used to temporarily permeabilize the plasma membrane of mammalian cells for the purpose of delivering materials to the cytosol. However, electroporation can be harmful to cells, while efficient mechanoporation strategies have not been scalable due to the use of narrow constrictions or needles which are susceptible to clogging. Here we report a method of mechanoporation in which cells were stretched and permeabilized by viscoelastic flow forces without surface contact. Inertio-elastic cell focusing aligned cells to the center of the device, avoiding direct contact with walls and enabling efficient (95%) intracellular delivery to over 200 million cells per minute. Functional biomolecules such as proteins, RNA, and ribonucleoprotein complexes were successfully delivered to Jurkat cells. Efficient intracellular delivery to HEK293T cells and primary activated T cells was also demonstrated. Contact-free mechanoporation using viscoelastic fluid forces appears to be feasible method for efficient and high throughput intracellular delivery of biomolecules to mammalian cells *ex vivo*.

---

Emerging cell and gene therapies promise to revolutionize oncology and other fields but face unprecedented challenges in manufacturing at clinical scales. One major obstacle has been the cost and complexity associated with efficiently delivering genetic materials and/or gene editing systems into very large numbers of cells *ex vivo* (hundreds of millions, up to trillions of cells for some applications)^1–3^. Broadly, there are three major *ex vivo* delivery methods currently in use for clinical-scale manufacturing: viral vectors, synthetic vectors, and high-throughput electroporation. Viral vectors such as lentivirus are unable to target a specific genetic locus, have a limited payload, and can be expensive to manufacture^4–6^. On the other hand, synthetic vector systems based on the formation of cationic/lipidic complexes can be unstable, inefficient, and/or cytotoxic^7, 8^. Unlike vector-based systems, reversible membrane disruption or ‘poration’ can permit the efficient delivery of many different classes of molecules into a wide variety of cell types^9^. Electroporation is a method using brief pulses of intense electric field to create temporary pores in the plasma membrane and move charged macromolecules into contact with the cell. However, electroporation is intrinsically biased towards the formation of numerous small pores (i.e. 2 nm) across a large fraction of the plasma membrane^10, 11^. To expand membrane pores for intracellular delivery, much higher E-field intensities and pulse durations are required that can cause DNA damage^12^, lipid peroxidation^13^, phenotypic changes and/or cell death^14, 15^. Scaled-up electroporation schemes face additional challenges associated with nonuniform E-fields, heating, electrolysis, pH changes, electrode corrosion, ionic contamination, and prolonged exposure of cells to low-conductance electroporation buffer^9^. Some of these issues with scale-up may be addressed using microfluidics,^16, 17^ however the intrinsic cytotoxicities associated with electroporation remain unresolved.

Lipid membranes may also be porated by applying one or more brief pulses of intense mechanical tensile stress, also called mechanoporation.^18^ Compared with electroporation, mechanoporation is thought to result in fewer and larger pores, since mechanopores tend to act as stress concentrations while electropores tend to relax the nearby potential.^19^ This would explain why mechanoporation can permit the efficient delivery of very large macromolecules or nanomaterials, can be well tolerated by many cell types when precisely controlled^18, 20–24^. However, highly controlled mechanoporation processes such as microinjection have proven difficult to scale. Techniques involving needles or some other surface physically contacting the membrane are intrinsically susceptible to clogging and/or fouling during prolonged use.^23, 25–28^ As an alternative, ‘non-contact’ mechanoporation using acoustic waves^29, 30^ or fluid shear stress^22, 31–34^ have been found to be much more scalable (millions of cells per minute) but at the expense of inefficient delivery and increased cell death. For example, the highest throughput mechanoporation method in the literature used microscale vortices to process up to 120 million cells per minute, but this highly nonuniform flow field resulted in only moderate delivery efficiency (less than 65% of cells) and significant cell death.^35, 36^ Theoretical, experimental, and computational studies have emphasized the importance of precisely controlling both the intensity and timescale of the applied membrane tension to avoid both subcritical cell stretching and catastrophic cell lysis.^19, 37–39^ Therefore, we investigated whether a scalable method could be devised of maintaining highly consistent mechanoporation conditions for all cells in a sample.

In this article we report a high throughput, non-contact method of mechanoporation which utilized viscoelastic flow phenomena within a microfluidic chip to maintain an approximately constant intensity and duration of applied membrane tension for all cells (Figure 1). Cells were suspended with the ‘cargo’ molecule to be delivered in a viscoelastic dilute polymer solution. Within the chip, cells were focused to the center of a microfluidic channel, preventing contact with walls and minimizing cell-to-cell variability in the residence time and peak stress. A contracting microchannel caused cells to be stretched by the relative motion of the surrounding viscoelastic fluid. The flow kinematics were characterized using experimental flow visualization and computational methods. The device and protocol were optimized, then used to successfully deliver mRNA and CRISPR-Ca9 ribonucleoprotein (RNP) complexes to Jurkat cells. Finally, intracellular delivery to HEK293T cells as well as primary T cells was evaluated, towards the goal of improving the safety, efficiency, and speed of *ex vivo* genetic manipulation for cell and gene therapy manufacturing.

**Figure 1.**
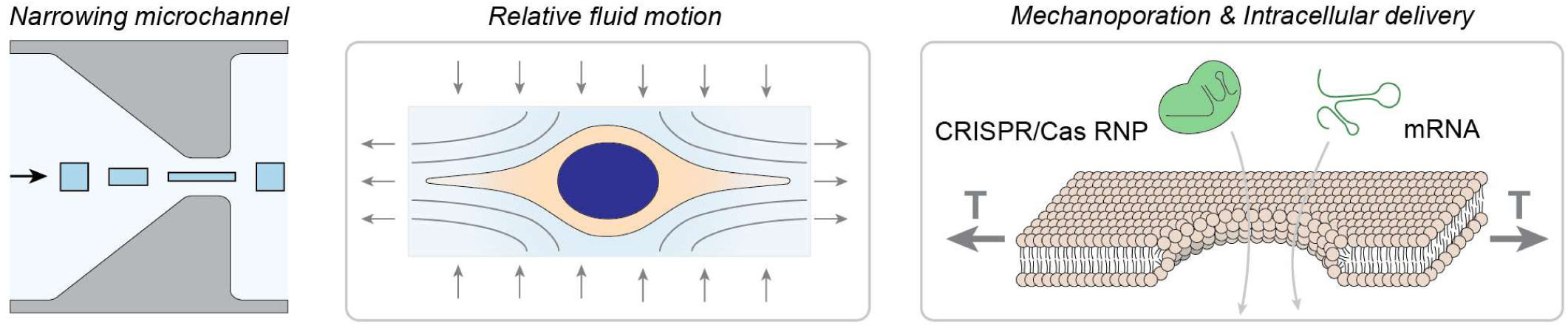
Conceptual schematic for high throughput continuous-flow transfection by flow forces. A fluid element that enters the center of a narrowing channel becomes elongated along the direction of travel (also known as an ‘extensional flow’). The plasma membrane of a cell in suspension is tensioned and permeabilized by inertial and drag forces from relative fluid motion, allowing intracellular delivery of biomolecules. The extensional viscosity of the solution is greatly increased by adding a biopolymer to make the solution viscoelastic, resulting in higher membrane tension at a given flow rate.

## Results

### Viscoelastic mechanoporation is feasible for efficient intracellular delivery of large biomolecules

A simple microfluidic contraction-expansion channel was initially fabricated from polydimethylsiloxane (PDMS) to test whether such a flow could generate sufficient membrane tension for mechanoporation of Jurkat cells, an available T cell line. This preliminary device consisted of a single straight microchannel 100 µm long, 45 µm wide, and 50 µm tall, that connected two much larger chambers (1.5 mm wide, 2 mm long, and 50 µm tall) by tapered sections (Figure S1). The channel was therefore at least three times wider than the cells, which were between 9 µm to 14 µm (Figure S2). Using this chip layout, we investigated the impact of several process parameters (polymer concentration, flow rate, channel geometry, and buffer composition) on metrics of delivery performance (delivery efficiency, cell viability, and the percentage of cells recovered after processing). 2,000 kDa Fluorescein isothiocyanate labeled dextran (‘FITC-dextran’) was used for optimization and characterization studies as an inert fluorescent dye of specified molecular size. For each trial, 50 µL of a delivery solution consisting of phosphate buffered saline (PBS) with 1.5 MDa hyaluronic acid (HA, a biocompatible polymer creating a viscoelastic solution), cells, and cargo molecule was pumped through the device at a fixed pressure.

Across tested all geometries and HA concentrations, the delivery efficiency increased with increasing flow rate, with delivery efficiencies exceeding 90% for some configurations (Figure S3-S5). In contrast, the delivery efficiency did not exceed 34% if the polymer was excluded. At higher flow rates, the viability decreased slightly while the cell recovery rate decreased substantially, suggesting substantial cell loss due to mechanical lysis. The addition of calcium ion in the transfection solution was also evaluated, since prior work has established the role of Ca^2+^ influx as an important signal for membrane repair.^40–42^ Indeed, we found a statistically significant increase in viability, from about 80% to about 90%, assessed 3 hours after transfection, when calcium was included in the transfection solution (Figure S6).

### High throughput cell focusing enables high throughput and consistent cell deformation

We hypothesized that upstream cell focusing would improve process consistency and cell recovery. We developed and integrated a custom cell focusing component that consisted of to two symmetrical outward-spiraling microchannels, each of which focused cells to an equilibrium position along of the outer (concave) wall of the spiral (Figure 2a-d). It has been shown previously that inertio-elastic flows in curved channels also focus cells vertically, i.e. halfway between the top and bottom of the channel.^43^ We quantified the focusing behavior of Jurkat cells suspended in PBS with a range of HA concentrations, flow rates, and channel heights (Figure 2d). All tested combinations of channel geometry and HA concentration were able to achieve efficient focusing at some range of optimal flow rates around 1 mL/min to 2 mL/min (Figure 2e and Figure S7). A channel height of 80 µm was selected for further study as it exhibited excellent focusing performance across the largest range of polymer concentrations and flow rates, particularly at higher flow rates.

**Figure 2.**
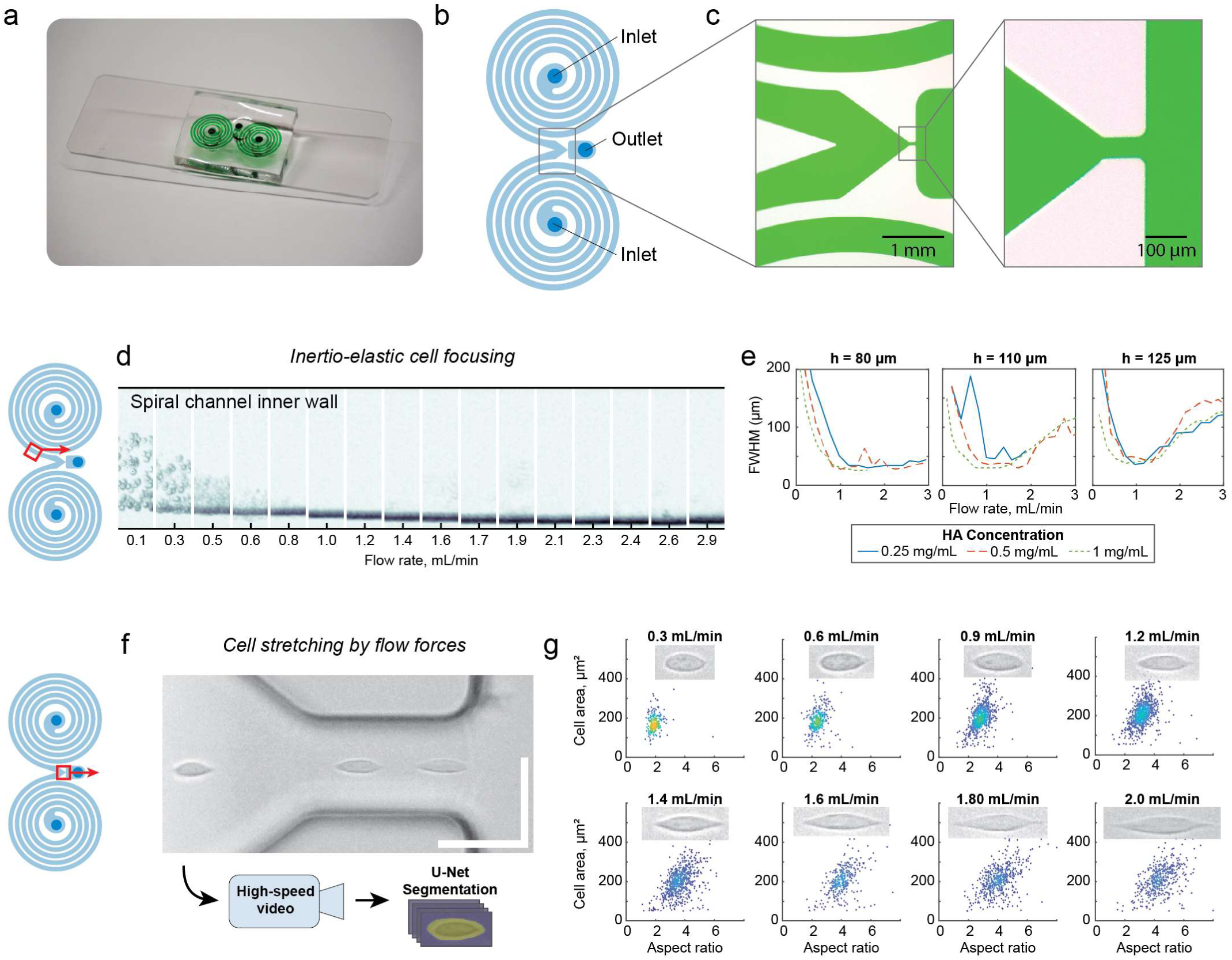
High throughput and uniform cell stretching without surface contact. (a) Image of epoxy-glass microfluidic chip with dye-filled channels. (b) chip layout and (c) microscope images showing the narrowing microchannel section where cells are stretched by flow forces. (d) Composited time-lapse images of one of the spiral channels show cells being focused to a narrow band near the outer (concave) channel wall. Indicated flow rate is the total flow through both inlets. Composite images are 500 μm tall. (e) Full-width half-maximum (FWHM) of lateral cell position distributions across the width of the composite images, for a range of channel heights and hyaluronic acid concentrations. Lower FWHM indicates tighter cell focusing. (f) High-speed video of cells in the narrowing channel were captured and segmented for quantitative shape analysis (scalebar indicates 50 μm). (g) Cell area and aspect ratios for several hundred cells at a range of flow rates, with representative inset images. Inset images are 16 μm tall.

Cell deformations within the extensional flow were imaged by high-speed video microscopy and quantified using a convolutional neural network (Figure 2f-g). As expected, cells were elongated along the flow direction and cell aspect ratio increased with increasing flow rate. Qualitatively, at lower strain rates cells remained rounded, while at higher strain rates cells were transiently pulled into an elongated spindle-like morphology, consistent with previous studies of transient cell deformation in pure extensional flows.^22, 32, 33^ Above 2 mL/min, cell imaging was degraded by motion blur due to very high flow speeds (>10 m/sec) and the minimum exposure time being limited to 0.3 µsec.

### Flow visualization and computational studies reveal contraction inlet vortices

The flow kinematics in the contracting channel were studied using experimental flow visualization and computational simulations. For a viscoelastic solution of 0.5 mg/mL HA in PBS, the pathlines of 2 µm beads revealed the onset of symmetric flow separation and upstream vortices at about 0.05 mL/min, followed by asymmetric vortex growth and oscillating instabilities above about 0.2 mL/min (Figure 3a). The upstream length of the vortices increased with flow rate, as did the frequency of oscillations (Figure 3b).

**Figure 3.**
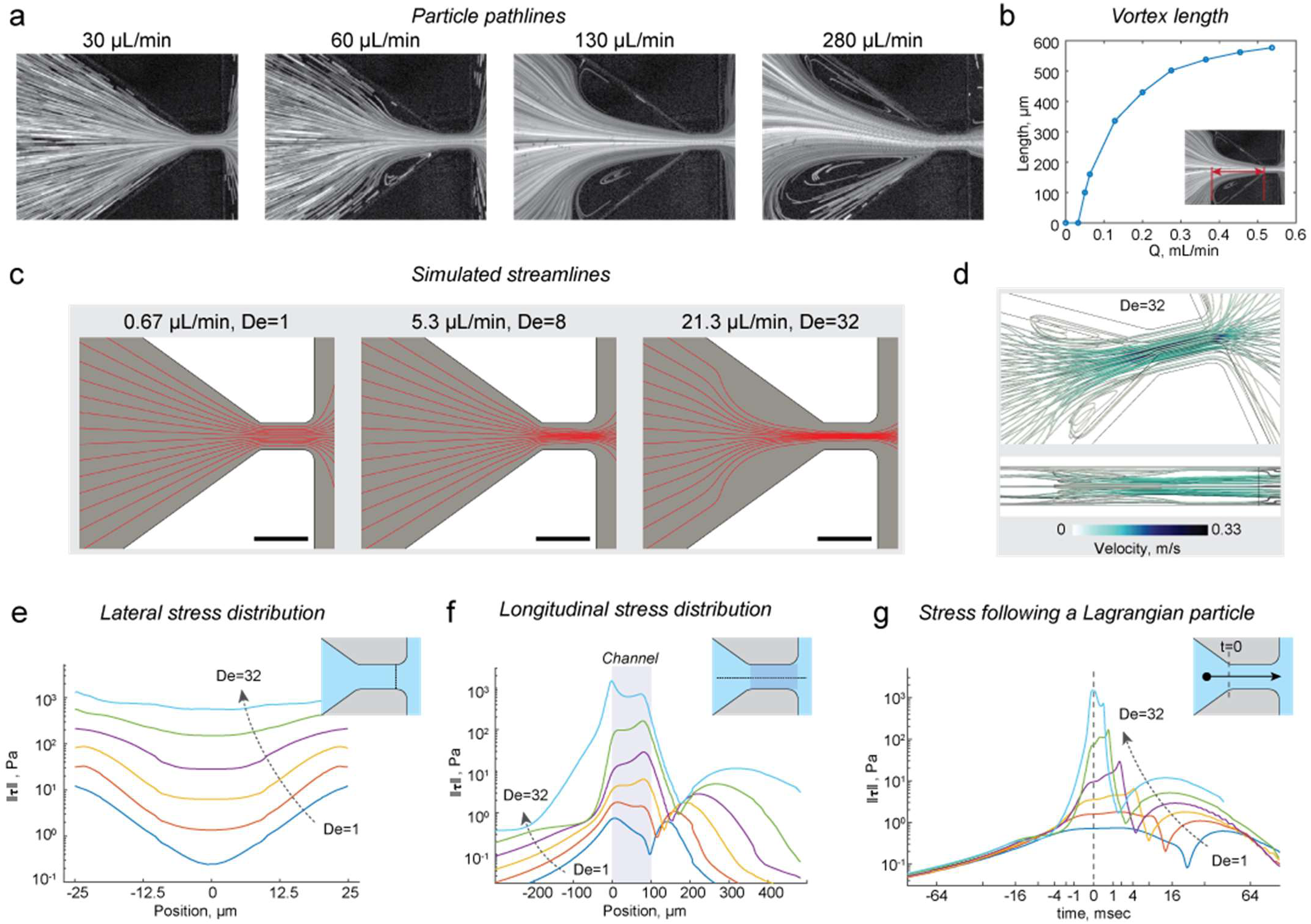
Experimental and computational characterization of the flow. (a) Pathlines show flow separation and vortices at increasing flow rates. (b) Vortex length as a function of flow rate. (c) Simulated streamlines of a viscoelastic FENE-CR fluid recapitulate vortices at 20 μL/min, Deborah number = 32. (d) Streamtubes colored by velocity show flow kinematics in 3D. (e-f) From simulations, line cuts of the spectral norm of the deviatoric stress (‖𝝉‖, described in text) for De = 1, 2, 4, 8, 16, and 32. (g) Simulated histories of ‖𝝉‖ for a Lagrangian particle which enters the constriction at t = 0. Scalebars in (a) and (c) indicate 100 μm.

Finite element simulations of the contracting flow were used to investigate the flow kinematics and internal stress. A fully 3D model of the microfluidic contraction was solved for a range of flow rates up to 0.0213 mL/min, corresponding with Deborah numbers (De) from 1 to 32 (discussed further in Supplemental Information, Figure 3c-d). We elected to use the characteristic Deborah number rather than the Weissenberg number, since the viscoelastic normal stress differences are not expected to be fully-developed^44^. The Deborah number given by 𝐷𝑒 = 𝜆𝛾̇ compares the characteristic deformation rate ̇ = 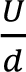 (with flow rate U and length scale 𝑑) with the fluid relaxation time 𝜆, an intrinsic property of the solution associated with the timescale of polymer rearrangement. The range of flow rates accessible to computational study were unfortunately limited to less than 1% of those required for mechanoporation (over 2 mL/min), due to fundamental challenges associated with finite element simulations of fast viscoelastic flows, which we discuss later. Nevertheless, simulations recapitulated the onset of flow separation at a similar critical flow rate to the experiment and supported several predictions about the stress distribution at high flow rates. To visualize how the stresses varied in a rotation-invariant way, we created plots of the spectral norm ‖𝝉‖ of the deviatoric stress tensor 𝝉 (i.e., the magnitude of the largest eigenvalue of the 𝝉). For a hypothetical small particle, ‖𝝉‖ may be considered analogous to the first principal stress. ‖𝝉‖ is relevant here because it is the extensional component of any given flow field that is responsible for cell stretching and mechanoporation^37^. We plotted the distribution of ‖𝝉‖ across the channel (Figure 3e) and along the channel centerline for increasing De (Figure 3f). To clarify how ‖𝝉‖ might vary over time for a hypothetical particle, we also calculated the stress history following the center streamline was calculated following a Lagrangian particle (Figure 3g). At low flow rates (De=1), drag from walls was dominant, and stresses along the channel centerline were much less than those at the walls. However, at higher flow rates, the contributions to stress from the flow contraction grew faster than contributions from wall drag, resulting in a more uniform lateral stress distribution. Also, while at De=1 the stresses associated with the flow contraction and expansion were comparable, at high flow rates the stresses associated with flow acceleration exceeded that of deceleration by over two orders of magnitude.

### Optimization of viscoelastic mechanoporation for high throughput and uniform intracellular delivery

We optimized the delivery efficiency, viability, cell recovery, and throughput performance of the device with integrated focusing (Figure 4a-h). The device was operated at pressures up to 7 bar, corresponding to flow rates up to 4 mL/min (Figure S8-S9). Compared with the contraction-only device, the cell recovery rate did not decrease as dramatically at the higher flow rates (Figure 4d), suggesting that cell lysis was greatly reduced by cell focusing. An overall yield was defined as the number of viable and dextran positive cells recovered as compared to the number of viable cells recovered in the unstretched control samples (Figure 4e). At the optimal operating pressure of 5 bar, the flow rate through the chip was about 2.7 mL/min. After 24 hours in culture, cell viability was slightly reduced but still above 90% for all samples and the cell number was about twofold higher, suggesting successful cell recovery and proliferation (Figure 4f-g and Figure S10). We also evaluated whether increasing cell concentration in the delivery solution could be used to increase throughput. At an operating pressure of 5 bar (nominally 2.7 mL/min) the delivery and viability did not decrease with increasing cell concentrations even at the highest tested concentration of 100 million cells per minute, supporting the feasibility of high throughput intracellular delivery of hundreds of millions of cells per minute (Figure 4h).

**Figure 4.**
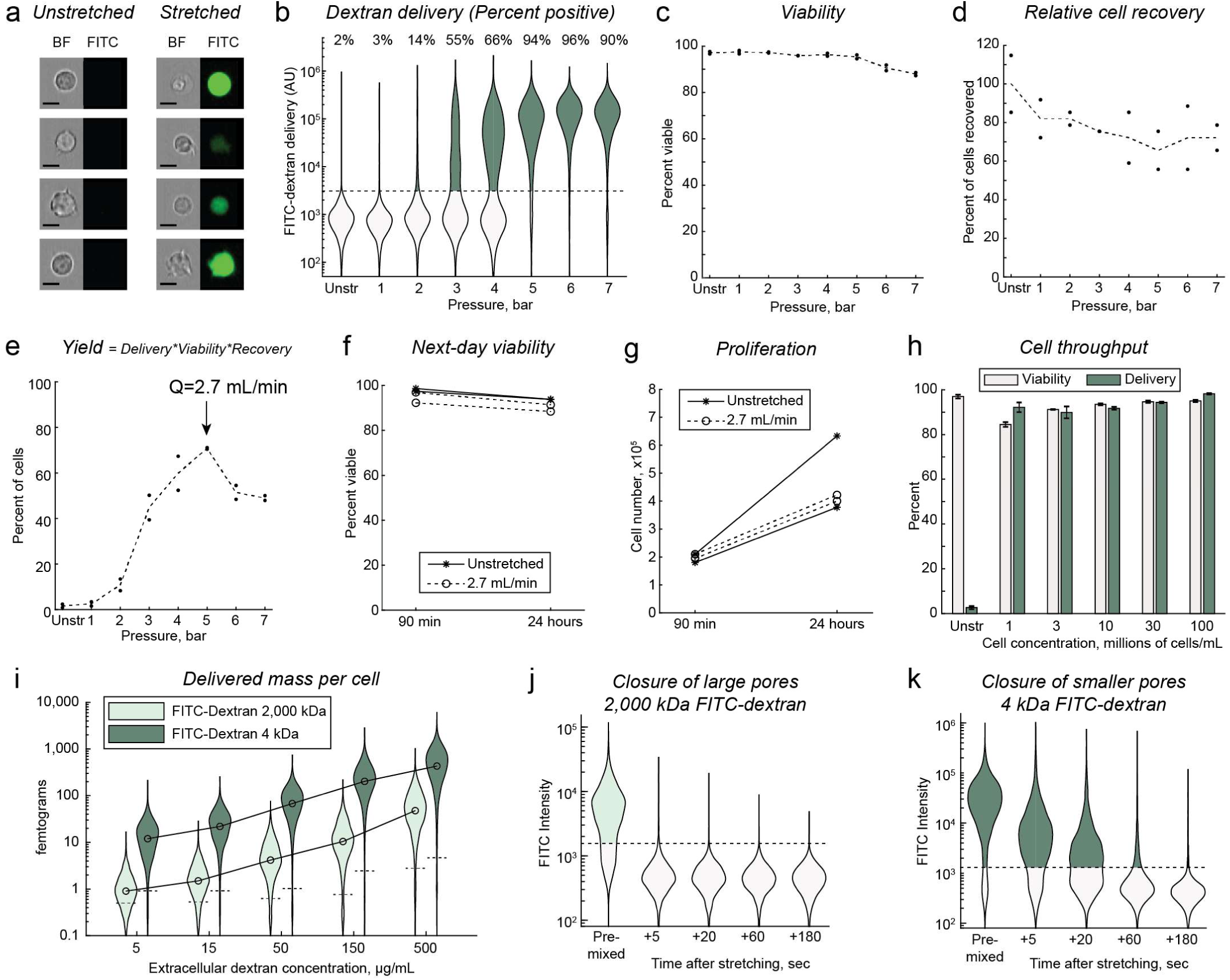
Optimization and characterization of mechanoporation for intracellular delivery. (a) Representative brightfield (BF) and fluorescence (FITC) images of Jurkat cells after viscoelastic mechanoporation and delivery of 70 kDa FITC-dextran at 2.7 mL/min. Images are padded to uniform size. Scalebars indicate 10 μm. (b) Distributions of intracellular FITC-dextran fluorescence of viable cells after processing cells with dextran through the chip at a range of pressures. ‘Percent positive’ indicates the percentage of viable cells that are brighter than an arbitrary cutoff (dotted line). ‘Unstr’ indicates ‘unstretched’, i.e., cells were incubated with dextran but not processed through the chip. (c) Cell viability and (d) number of recovered cells relative to unstretched samples, about 90 minutes after processing (n=2). (e) Yield is the number of viable and dextran positive cells recovered as compared to the number of viable cells recovered in the unstretched control samples. (f) Viability and (g) total cell number following 90 minutes or 24 hours of culture after processing. (h) Viability and dextran delivery for samples where the cell concentration was increased (n=3). (i) Distributions in the total mass of delivered material per cell, for large and small dextrans at a range of extracellular dextran concentrations. (j-k) Distributions in the amount of delivered dextran per cell when the dextran is added following a delay after processing, rather than premixed with the solution before stretching as usual. Dotted lines in (ik) indicate the arbitrary cutoff for positive delivery, i.e., 98th percentile of unstretched controls (not shown).

Quantitative flow cytometry was used to measure the mass amount of delivered material for both small and large biomolecules as a function of extracellular concentration (Figure 4i, calibration data Figure S11). Across all extracellular concentrations, the average amount of 4 kDa dextran delivered per cell was roughly 10-fold higher than for 2,000 kDa dextran. If a constant cell diameter of 12 µm was assumed (based on the average cell size, Figure S2), the average intracellular concentration of 4 kDa dextran fell within a factor of two of the extracellular concentrations across all tests (Figure S12). To assess how long the cell membrane remained permeable after mechanoporation, cells were processed through the chip without dextran in the solution, and then either small (4kDa) or large (2,000 kDa) FITC-dextran was spiked into the solution after a time delay of up to three minutes (Figure 4j-k). Some uptake of 4 kDa dextran was observed up to 20 seconds after processing, albeit greatly reduced, while uptake of 2,000 kDa dextran was undetectable even if added just five seconds after mechanoporation.

To verify that viscoelastic mechanoporation was due to the viscoelastic fluid properties rather than some biochemical effect specific to HA, viscoelastic solutions were prepared instead using 2,000 kDa polyethylene oxide (PEO) at concentrations of 1 mg/mL and 0.2 mg/mL in PBS. As expected, delivery of 70 kDa FITC-Dextran to Jurkat cells increased with increasing operating pressures, with efficient (>85%) delivery observed at some operating pressure for each tested PEO concentration (Figure S13).

### Viscoelastic mechanoporation can deliver protein, mRNA, and ribonucleoprotein complexes to Jurkat cells

We next evaluated whether viscoelastic mechanoporation could deliver broader categories of biomolecules. Using the optimized protocol, we found the delivery efficiency of FITC-tagged albumin to Jurkat cells was about 90% (Figure S14). Delivery of mRNA encoding enhanced green fluorescent protein (eGFP) was evaluated for a range of extracellular mRNA concentrations. Viability and eGFP expression were evaluated 24 hours after processing. As expected, the distribution in cell expression of eGFP increased with increasing extracellular concentrations of mRNA (Figure 5a). Following delivery of 100 µg/mL mRNA, 89% of viable cells were eGFP positive compared to 2% of cells that were incubated with the mRNA but not processed through the chip. Viability decreased with increasing amount of delivered mRNA, from 95% when no mRNA was delivered to 74% for 100 µg/mL, suggesting some potential cytotoxicity associated with the delivered material (Figure S15).

**Figure 5.**
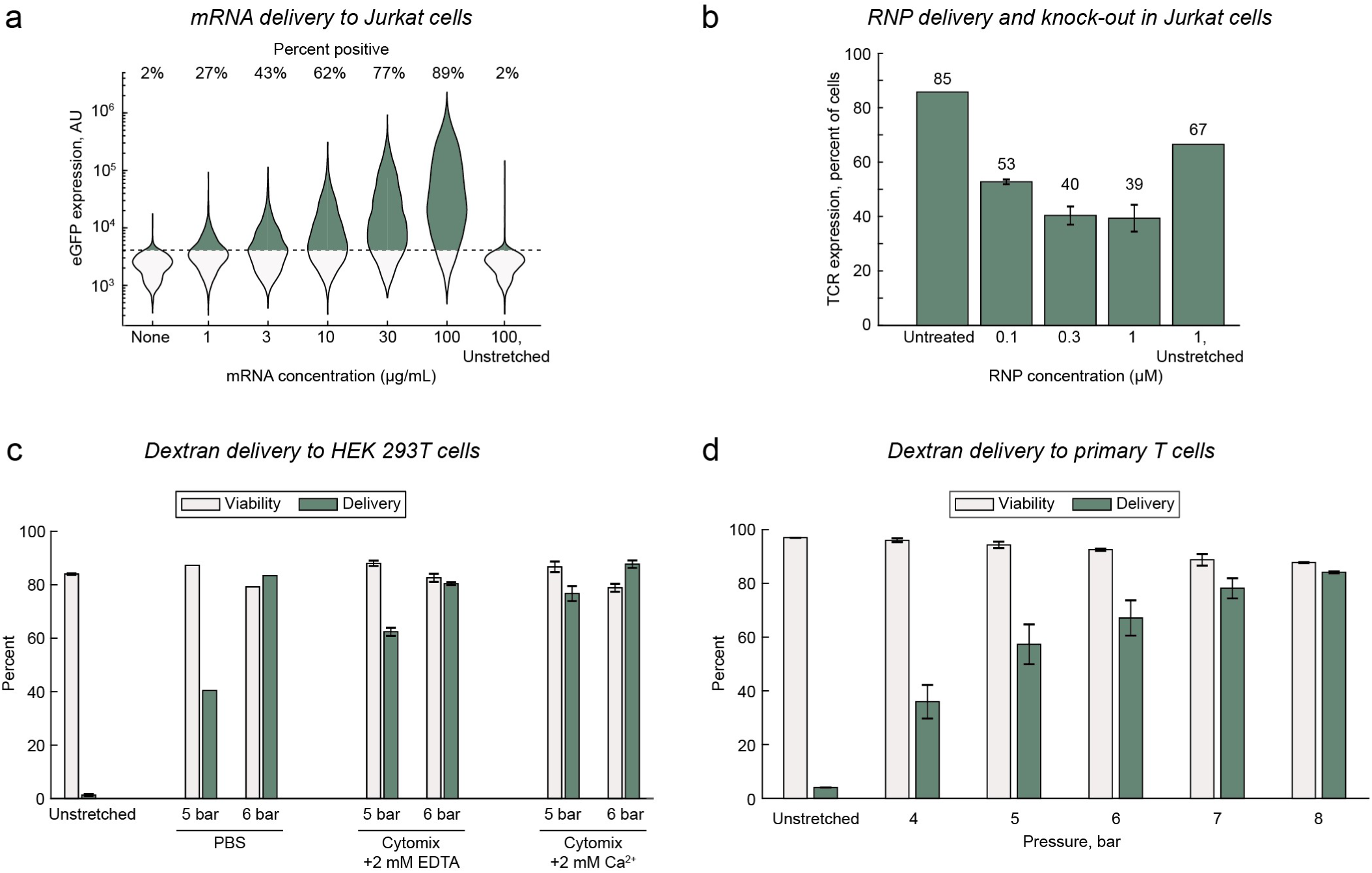
Generalization to different cargo molecules and cell types. (a) eGFP expression 24 hours after mRNA transfection to Jurkat cells for a range of extracellular mRNA concentrations. (b) T cell receptor (TCR) expression in Jurkat cells 2 days after transfection of Cas9 RNP complexes designed to knock out the TCR, for a range of RNP concentrations (n=3 for experimental conditions, n=1 for controls). (c) Viability and delivery of 70 kDa FITC-dextran to HEK293T cells, 24 hours after processing, for three delivery buffer compositions and 2 different chip pressures (Unstretched, n=2; PBS, n=1; Cytomix buffers, n=3). (d) Viability and delivery of 70 kDa FITC-dextran to primary activated T cells, about 90 minutes after processing, for increasing chip pressures (n=2).

RNP delivery for gene editing was evaluated in Jurkat cells using a synthetic crRNA sequence previously optimized to knock out the T cell receptor (TCR, Table S1).^45^ RNPs were complexed *in vitro* at a ratio of 1:2:2 Cas9:crRNA:tracrRNA, then added to the delivery solution at a final Cas9 concentration between 0.1 µM to 1 µM. RNP were delivered using the optimal protocol and cultured for two days. TCR expression was assayed by immunofluorescence staining and flow cytometry. As expected, the percentage of cells expressing detectable levels of TCR decreased with increasing RNP concentrations, from about 85% of untreated cells to less than 40% of cells treated with 0.3 µM or 1 µM RNP, indicating an average knockout efficiency of about 53% (Figure 5b). The viability of cells transfected with RNP decreased to between 66% and 73% after two days, although without a dose-dependent trend, which suggested some degree of cell damage associated with RNP delivery by mechanoporation (Figure S16).

### Viscoelastic mechanoporation of HEK cells and primary T cells

We next evaluated whether viscoelastic mechanoporation could be used to deliver molecules to other cell types, such as adherent cells or primary cells. We first optimized a protocol for delivery of 70 kDa FITC-dextran to HEK293T cells, an available adherent cell line. Cells were initially processed in 0.5 mg/mL HA in PBS at a range of operating pressures. Efficient delivery was observed at the highest operating pressures (>90%), although viability after about 90 minutes was relatively low at below 50% (Figure S17). We hypothesized that viability may be improved by using a ‘cytoplasmic’ buffer for the delivery solution which instead approximated the ionic composition of the cytosol, such as is used for nucleofection^46^. We were also interested to evaluate whether extracellular calcium would be helpful or harmful to cell recovery, as Ca^2+^ influx is a required signal for rapid membrane repair but can also trigger apoptosis. Overall, the cytoplasmic buffer with supplemented calcium was found to be best, resulting in viability after 24 hours similar to that of the untreated controls, and delivery efficiency of about 88% (Figure 5c). Finally, we evaluated delivery of 70 kDa FITC-Dextran to primary activated and expanded T cells 7 days after isolation, at a range of operating pressures and HA concentrations (Figure S18). Ultimately, the highest tested HA concentration of 2 mg/mL was best, with an average delivery efficiency of 84% and viability of 85% at the highest tested HA concentration of 2 mg/mL, and operating pressure of 8 bar (Figure 5d).

## Discussion

In this work, we developed a high throughput method of applying consistent mechanoporation conditions to cells without surface contact. Cells were stretched and permeabilized using a viscoelastic flow in a contracting microchannel. Contact between cells and walls was actively prevented by a custom high throughput cell focusing component, which required the use of rigid prototyping materials. Efficient intracellular delivery was accomplished for suspension (Jurkat) and adherent (HEK293T) cell lines as well as primary T cells, supporting the feasibility of this method as a general approach for *ex vivo* delivery of biomolecules. Delivery and expression of mRNA was observed at extracellular concentrations similar to those typically used for electroporation (30 µg/mL – 100 µg/mL). Likewise, efficient delivery was observed with high cell concentrations (10^7^ – 10^8^ cells/mL). Altogether, viscoelastic mechanoporation seems to be a feasible approach for efficient intracellular delivery at throughputs of several hundreds of millions of cells per minute, per microchannel.

This approach should be compared to other recent developments in mechanoporation technologies, especially other microfluidic methods involving fluid forces. Cells have been porated by intense flow fields including cross-slot flows^34^, T junctions^22, 32^, vortices^35, 47^, and nebulizers^29^. Cells have also been observed to deform in highly viscous and/or shear-thinning solutions, and this was recently explored for improving delivery during microfluidic cell squeezing.^26, 48^ The important differences between viscoelastic mechanoporation and these methods are the process uniformity, throughput, and delivery performance.

Viscoelastic mechanoporation was capable of very efficient and uniform delivery (e.g., 95% efficiency for Jurkat cells) at a throughput that is much higher than the previous throughput record for mechanoporation, using micro-vortices (a highly nonuniform process that was much less efficient)^35, 36^, and at least two orders of magnitude faster than all other methods. Prior flow-based mechanoporation strategies also could not keep both the magnitude and the duration of membrane tension constant for all cells. This was because the flow conditions were either inhomogeneous (i.e., spatially varying, with cells in different positions experiencing different loads) or singular (i.e., cells were hydrodynamically trapped for uncontrolled amounts of time at a stagnation point, such as a junction or vortex where the net flow is zero). Relatedly, none of these methods were compatible with high cell densities (e.g., 10^8^ cells/mL) due to issues with clogging or the fact that only one cell should occupy the stagnation point at a time. In this work, clogging and uniformity issues were avoided by focusing cells to the center of a channel that was several times wider than the cells. The favorable performance and very high throughput of viscoelastic mechanoporation were enabled by leveraging intrinsic viscoelastic properties of dilute polymer solutions to achieve sufficient membrane tensions in a continuous-flow system.

The computational studies were limited to relatively low flow rates due to the so-called High Weissenberg Number Problem, a numerical stability issue that is well-known within computational rheology which makes it challenging to perform finite-element simulations of fast viscoelastic flows.^49^ As a result, virtually all published computational results of viscoelastic contracting flows have been 2D and/or neglected fluid inertia. Our studies, the longest of which took over three hundred thousand iterations and six months to solve, have provided (to our knowledge) the first solutions for a fully-3D contracting viscoelastic flow with appreciable inertia. At higher flow rates an unsteady flow condition was observed experimentally, which should be avoided to further improve the mechanoporation process uniformity.

The possibility of delivering the polymer into the cell was not investigated. Hyaluronic acid is readily digested by intracellular enzymes, but it can also be recognized as a biochemical signal. We showed that the method requires a viscoelastic solution but is not dependent on a specific chemical structure. This will permit material selection and optimization for different cell types as necessary.

Altogether, this work demonstrated that viscoelastic mechanoporation is feasible for high throughput delivery of biomolecules into mammalian cells *ex vivo*. Further optimization of the device geometry will mitigate inertio-elastic instabilities, while selection of different viscoelastic polymers will allow cell- or process-specific optimization for diverse biomedical applications. We expect these studies to inspire and enable further development of scalable intracellular delivery strategies for cell and gene therapy manufacturing.

## Methods

### Rigid microfluidic device fabrication and fixturing

Devices were microfabricated from molded epoxy bonded to glass. Master molds were fabricated by photolithography of SU-8 silicon wafers, and patterns transferred to a poly(dimethyl) siloxane replica mold used for epoxy casting, as described in detail elsewhere.^50^ 1.5 cm segments of 1/16^th^ inch ID polyethylene tubing were integrated for the device inlets and outlet during epoxy casting.

### Preparation of viscoelastic solutions

Stock solutions of 4 mg/mL HA were typically prepared by dissolving 1.6 MDa sodium hyaluronate (HA15, Lifecore Biomedical) in phosphate buffered saline (PBS) overnight with gentle rocking at room temperature. HA solutions were stored away from light at 4C and used within 2 weeks. Stock solutions of 5 mg/mL polyethylene oxide, 2MDa (PEO, Sigma-Aldrich) were prepared and stored similarly.

### Cell segmentation for automated cell deformation analysis

High-speed videos were acquired of cells passing through the contracting microchannel using brightfield phase contrast microscopy with a 40x objective. Videos were subsampled as necessary to ensure any individual cell did not appear more than once. Altogether about 48,000 frames were acquired across the eight different tested flow rates. Training data was generated by randomly selecting a subset of 3,000 frames and manually drawing individual cell outlines in ImageJ. The standard U-Net classifier was used, with outputs weighted by the prevalence of cell-pixels versus background-pixels in the training data.^51^

### Delivery experiments

Except where stated otherwise, cells were resuspended at concentrations between 3 million cells/mL to 10 million cells/mL in a solution of phosphate-buffered saline (PBS) containing the cargo molecule (e.g., 0.2 mg/mL 70 kDa FITC-dextran), 1.0 mg/mL HA and 0.2 mM CaCl2. The cell suspension was strained with a 40 µm nylon mesh prior to processing through the chip. The ‘cytoplasmic’ buffer used for HEK293T cells consisted of 100 mM KH2PO4, 15 mM NaHCO3, 12 mM MgCl2 × 6H2O, 8 mM ATP, and 2 mM glucose, to which with either 2 mM CaCl2 or 2 mM Ethylenediaminetetraacetic acid (EDTA) were added. For each experimental condition, 100 µL of the mixture was pipetted into each of two custom 3D printed reservoirs that attached to the inlet tubing by barbed Leuer fittings. The chip was primed with the sample solution by attaching an airtight pneumatic manifold to the tops of both reservoirs and using gentle pressure via a hand-operated syringe. Then, the inlet was pressurized by opening a pneumatic valve to a regulated pressure source. In this way, 200 µL samples were typically processed within about 5 seconds. The sample exiting the outlet tubing was collected and diluted immediately in room temperature culture medium (for Jurkat cells and T cells, RPMI 1640 supplemented with 10% fetal bovine serum and 100 u/mL pen-strep; for HEK293T cells, DMEM supplemented similarly). Cells were maintained at room temperature for up to one hour before being washed twice in fresh culture medium. Control ‘unstretched’ cell samples were suspended in the complete delivery solution and handled identically except they were not pumped through the chip. The delivery efficiency was defined as the percentage of viable (i.e., PI-negative) cells with a higher FITC signal than the substantial majority (e.g., 97%-98%, manually gated) of viable cells in an ‘unstretched’ control sample, for which cells were incubated in the delivery solution containing the dye but not processed through the chip. This permitted endocytotic uptake or cell surface binding to be distinguished from mechanoporation and intracellular delivery.

### mRNA and Cas9 RNP transfection

For mRNA transfection, we used an mRNA encoding enhanced green fluorescent protein (eGFP) with ARCA cap modifications (Apexbio Technologies, Fisher Scientific #50-199-8310). For Cas9 RNP complex transfection, 24 µL of 60 µM crRNA (Table S1) was hybridized with 12 µL of 120 µM atto550-tagged Atr-R™ tracrRNA (Integrated DNA technologies) in Tris-buffered saline by warming to 80 C for 5 minutes, then cooling to 4 C for 15 minutes. The hybridized RNAs were then mixed with 25 µL of 30 µM (5 mg/mL) recombinant SpCas9 protein (Sigma-Aldrich #Cas9PROT) for 30 minutes at 4 C, resulting in a molar ratio of 2:1:1 Cas9:crRNA:tracrRNA. T cell receptor (TCR) expression was quantified with anti-TCR labeled antibody (BioLegend #306706) and imaging flow cytometry.

### Isolation and culture of primary immune cells

Human peripheral blood mononuclear cells were isolated from whole blood by density centrifugation with Ficoll-Paque, and CD3 T cells were isolated from the PBMC fraction by incubation with anti-CD3/anti-CD28 magnetic beads following manufacturer’s instructions (Dynabeads, Thermo Fisher Scientific #11161D). Activated T cells were subcultured on day +3 and when cell density exceeded 1e6 cells/mL. Delivery experiments were uniformly conducted on day +7 after isolation.

### Flow visualization studies

Polystyrene tracer particles (2 µm, Stokes number < 0.001) were dispersed at 0.05% w/v in 0.5 mg/mL HA in PBS and imaged with a high-speed camera at 64,000 frames per second. Particle pathlines were generated by projecting the pixelwise standard deviation through several hundred frames.

### Computational fluid dynamics simulations

We used RheoTool, a package extending the capabilities OpenFOAM with stabilized numerical solvers and custom high resolution schemes optimized for simulations of viscoelastic flows.^52^ A high-resolution, fully-swept mesh was generated, consisting of about 2 million hexahedral elements, that followed the experimental geometry except that the channel height was set to 50 µm rather than 80 µm (**Figure S9**). The viscoelastic rheological parameters 𝜆 = 0.0125 sec (relaxation time) and 𝜂_0_ = 5 mPa · sec (zero-strain viscosity) were selected based on published rheological measurements of aqueous solutions of 1.5 MDa hyaluronic acid with concentrations in the range of 1 mg/mL to 2 mg/mL.^53–55^ Together with a characteristic channel shear length scale of 𝑑 = 25 µ𝑚 and fluid density of 𝜌 = 1000 kg/m^3^, these provided a characteristic Elasticity number 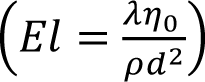 of exactly 100. The Elasticity number describes the ratio of elastic to inertial forces and is pd^2^ independent of the flow rate. The FENE-CR constitutive model was used, with the solvent and polymer contributions to the viscosity selected as ηs = 1 mPa*s and ηp = 4 mPa*s, respectively, and the chain length factor was set to 1,000, which is considered large for a FENE-type model (i.e. the simulated chains are long) yet compared to the biophysical chain length of 1.6 MDa hyaluronic acid of roughly 4,000 monomers per molecule.^56^

### Flow cytometer calibration for quantitative flow cytometry

The Amnis ImageStream MkII performs multispectral imaging using multiple laser excitation and a dichroic filter stack to perform spectral decomposition onto a single CCD sensor. For green-emitting fluorochromes, emission is collected from 505 nm to 560 nm with a band peak at 533 nm. For FITC excitation, a 488 nm laser is used. The Ultra Rainbow Calibration bead kit (Spherotech URCP-38-2k) consists of 3.8 µm polystyrene particles which contain embedded six different fluorochromes that align to the most common flow cytometry excitation and emission filter sets. The kit consists of five premixed populations of beads that are embedded with increasing amounts of the dye mixture. First, we determined the linearity of the fluorescence intensity signal in the FITC channel of the flow cytometer across the sensor dynamic range by comparing fluorescence intensities of each peak with the population MEFs (molecules of equivalent fluorochrome) as provided by the manufacturer (**Figure S11a-b**). These results indicated adequate linear response for reads between 1e3 and 2e5 AU. These data also provided the relative numbers of each bead population and the overall population average MEF per bead of 17,277. Next, we determined the equivalent MEF for FITC-dextran species used for delivery by comparing the fluorescence of the FITC-Dextran species to the calibration beads using a spectrophotometer (SpectraMax iD3). Excitation and emission wavelengths of 485 nm and 535 nm, respectively, were selected to match the flow cytometer FITC channel excitation and emission band peaks (488 nm and 533 nm). The bead concentration was measured by a Nexcelom cell counter. The resulting measurements provided equivalences of 1 MEF = 3.5 attograms of 2,000 kDa FITC-Dextran, or 5.4 attograms of 4 kDa FITC-Dextran. These equivalences, together with the calibration factors for fluorescence signal across increasing flow cytometer laser powers, allowed the flow cytometer fluorescence intensities to be converted to femtograms of FITC-dextran (Figure S11c).

## Acknowledgments

This work was supported by the National Institute of Allergy and Infectious Disease (K99 AI167063), the National Cancer Institute (R01 CA255602), and the National Science Foundation Engineering Research Center on Advanced Technologies for the Preservation of Biological Systems (#1941543). We thank Carlie Rein for assistance with microdevice fabrication, Shannon Stott for microscopy resources, and Jon Edd, Avanish Mishra, Kaustav Gopinathan, for helpful discussion.

## Author Contributions

D.S. and M.T. conceptualized the work, reviewed results, and reviewed the manuscript. D.S. designed and performed experiments and simulations, analyzed the data, and wrote the manuscript.

## Supplementary Information

**Figure S1.**
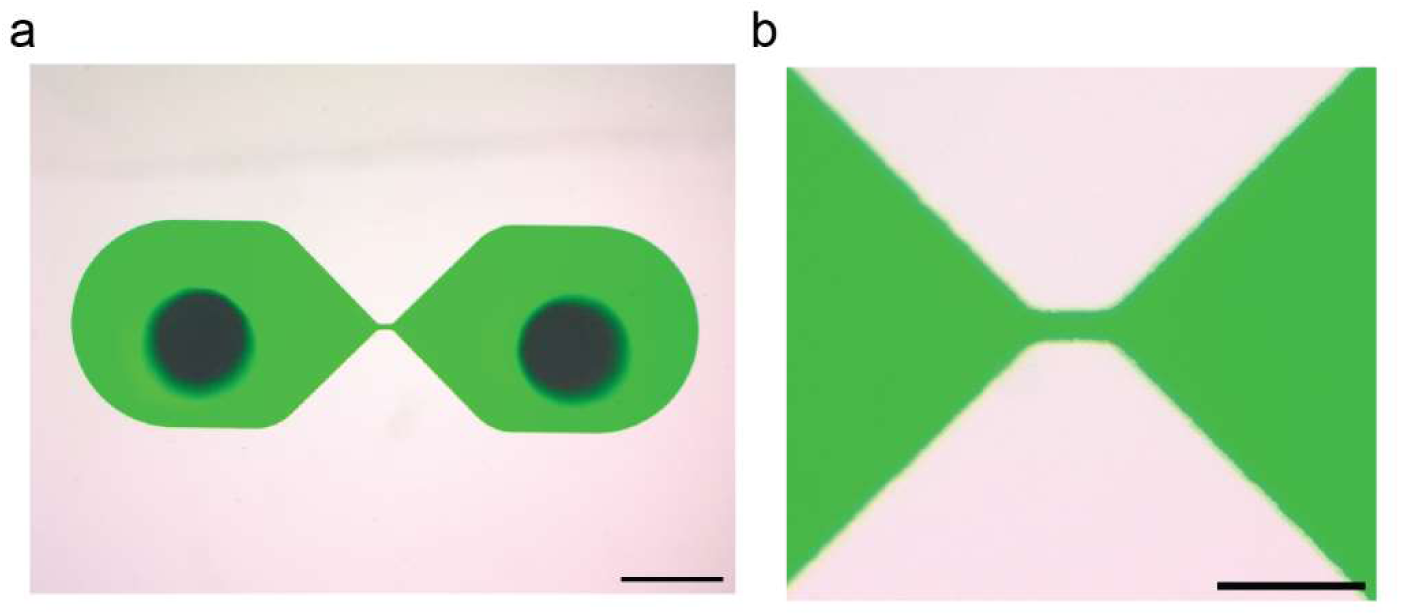
Images of initial microfluidic device without upstream cell focusing, visualized with green dye. Scalebars indicate (a) 1 mm and (b) 250 μm

**Figure S2.**
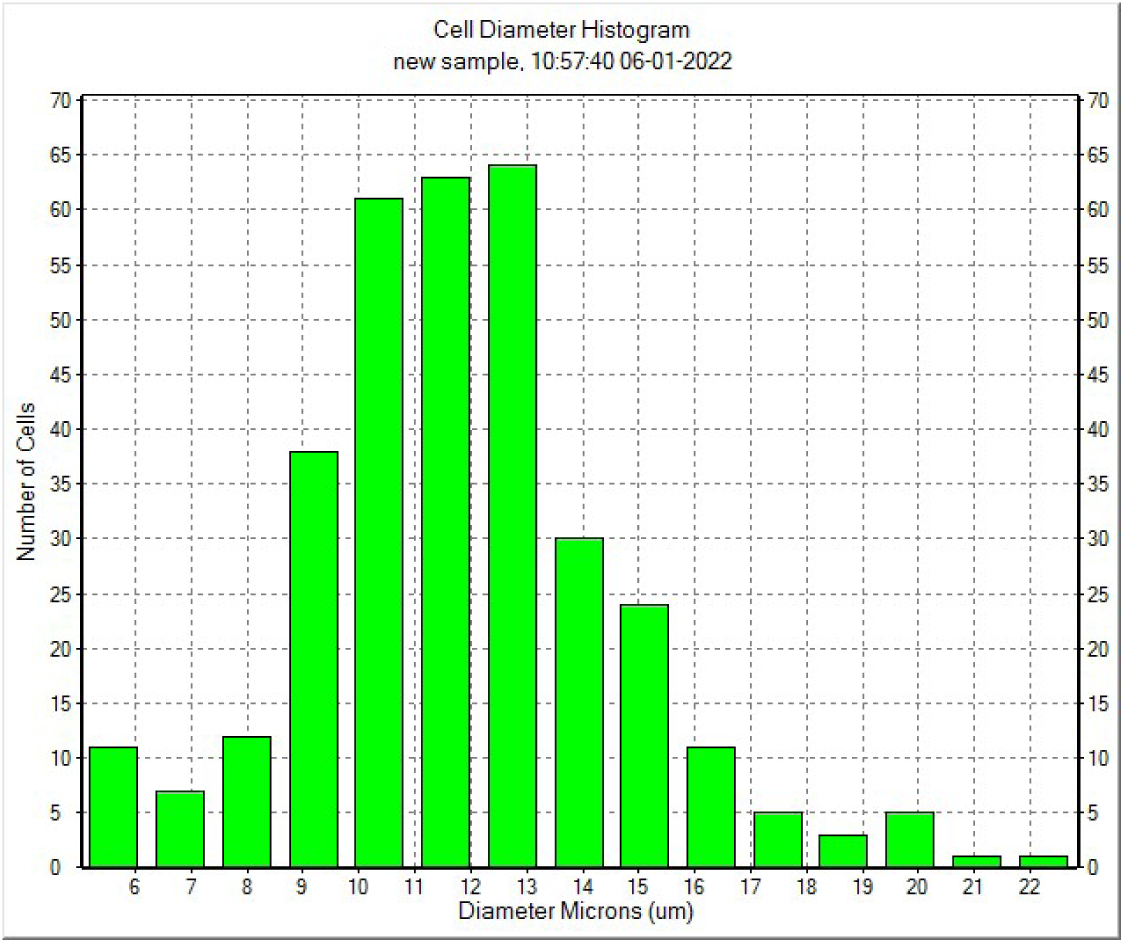
Representative distribution of Jurkat cell size.

**Figure S3.**
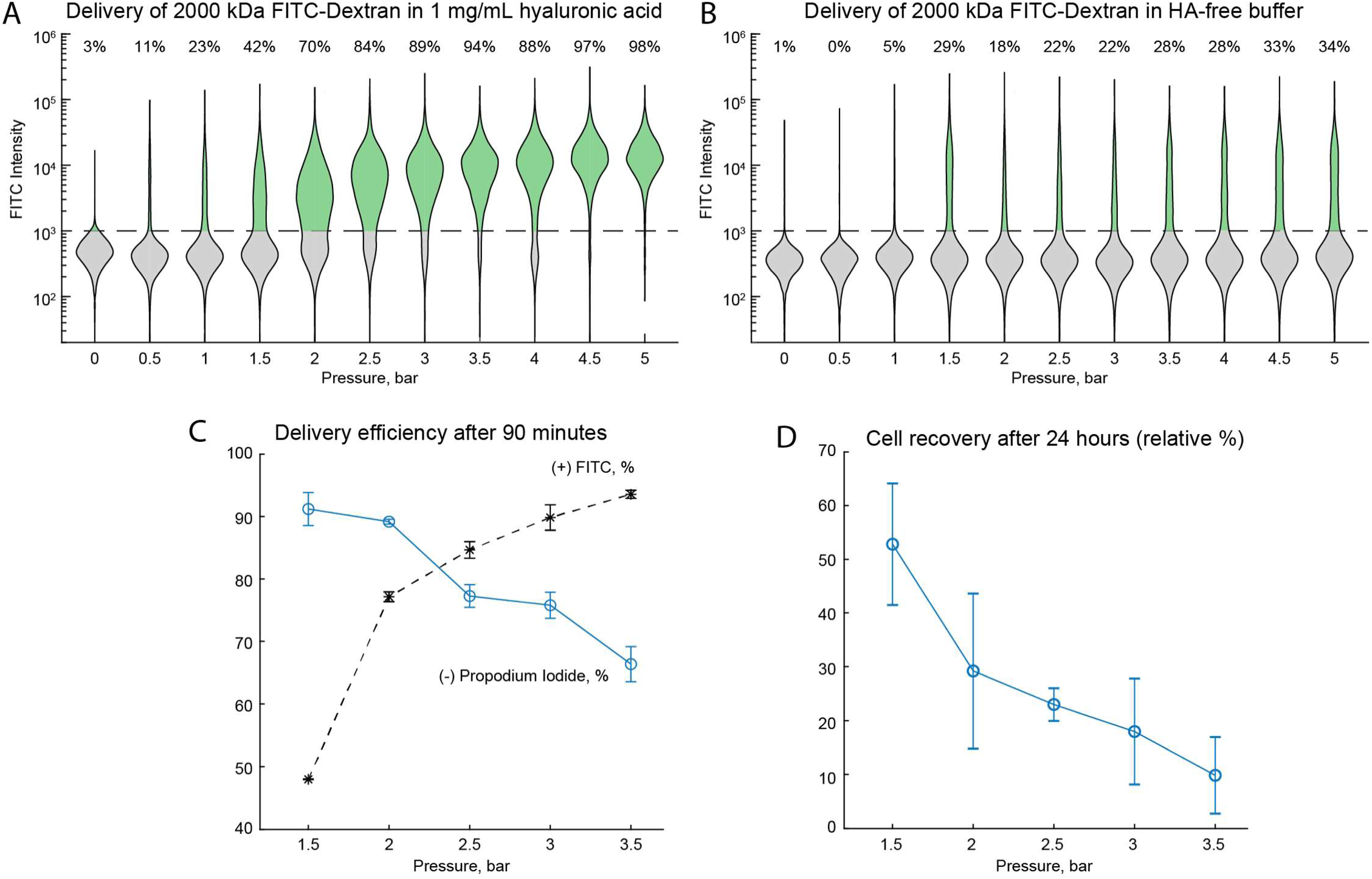
Transfection performance in the channel-only device. Distributions of FITC intensities for viable cells after processing in the channel-only device (Figure S1) at increasing driving pressure with a transfection solution containing (a) 1 mg/mL HA and (b) no HA. ‘0 bar’ indicates unstretched controls Percentages indicate delivery efficiency. (c) Delivery efficiency and viability 90 minutes after stretching in 1 mg/mL HA for a reduced range of pressures (n=3). (d) Total cell recovery relative to unstretched controls, following overnight culture.

**Figure S4.**
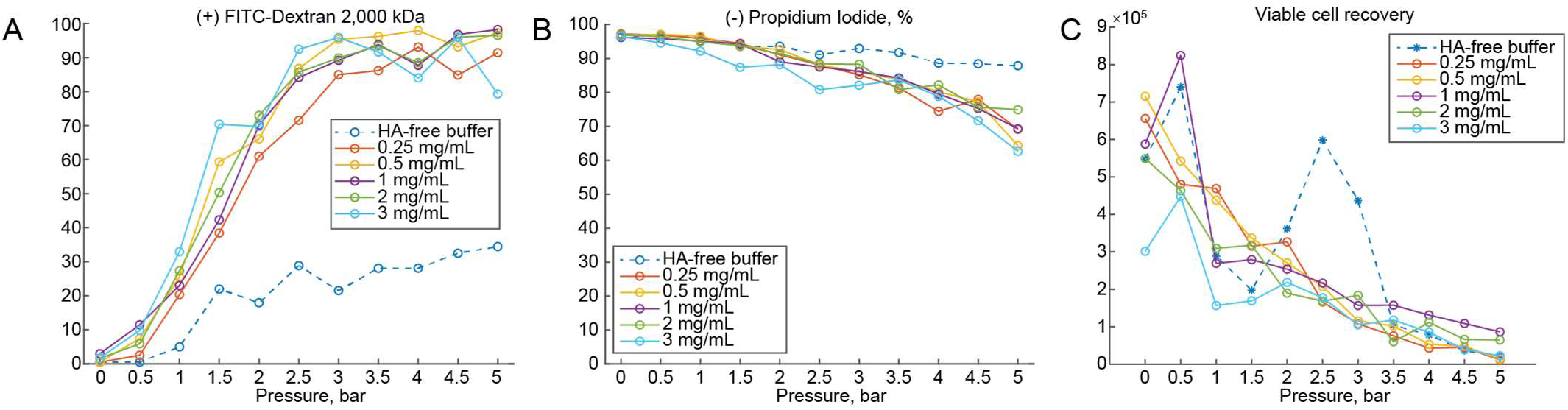
Optimization of transfection buffer in the channel-only device. Jurkat cells were suspended in transfection buffer of PBS with increasing amounts of HA and 0.3 mg/mL FITC-Dextran 70 kDa, and pumped through device at fixed pressures between 0.5 bar and 5 bar. (a) Delivery efficiency, (b) viability, and (c) total number of viable cells recovered were evaluated about 90 minutes after stretching. ‘0 bar’ indicates corresponding unstretched controls.

**Figure S5.**
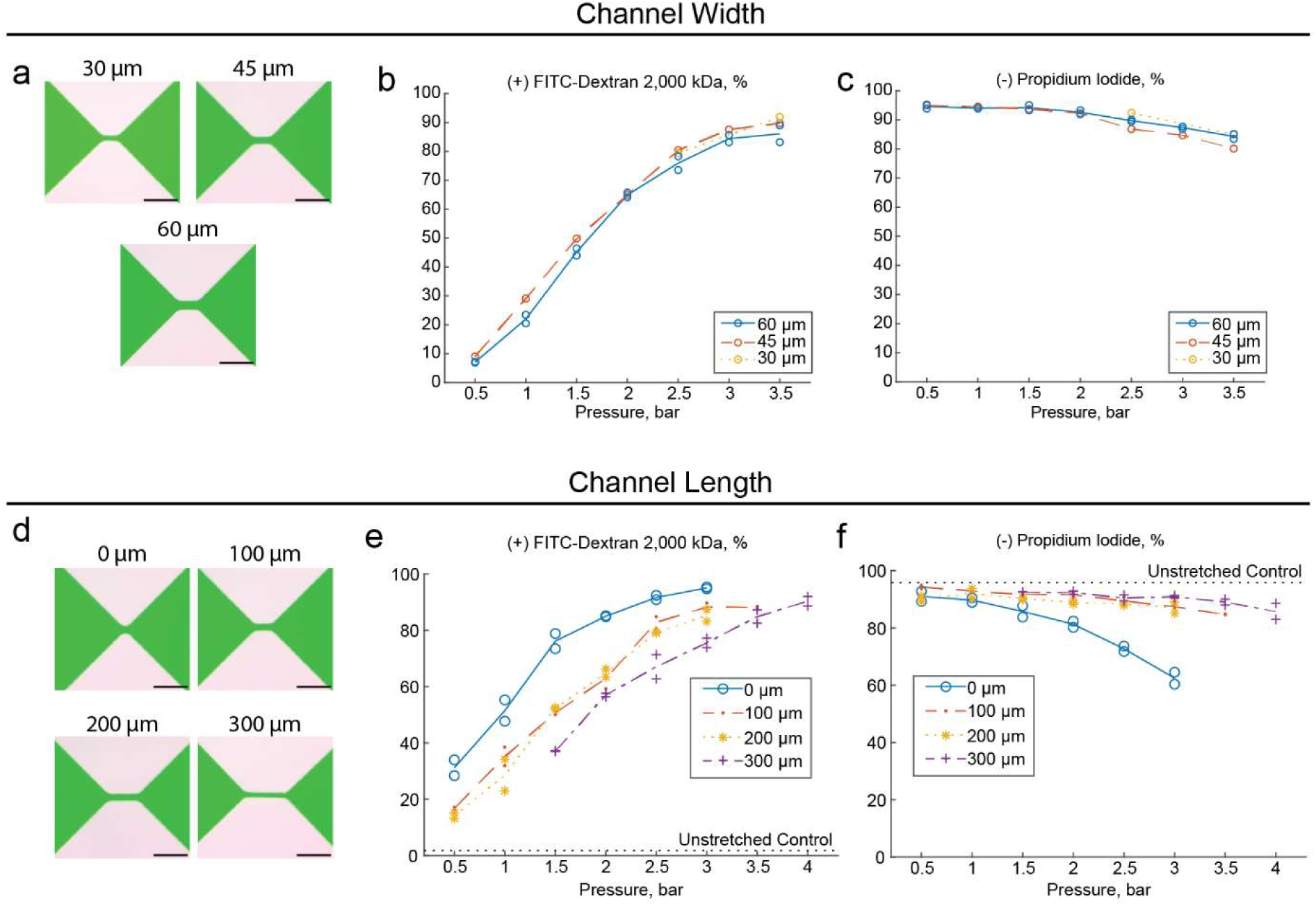
Mechanoporation performance for differing channel widths and lengths. Jurkat cells were suspended in transfection buffer of PBS with 2 mg/mL HA and 0.3 mg/mL FITC-Dextran 70 kDa, and pumped through devices of different channel widths at fixed pressures up to 5 bar. (a) Images of devices with channel widths 30 μm, 45 μm, or 60 μm wide (length was fixed at 100 μm). (b) Delivery efficiency and (c) viability 24 hours after cell stretching. (d) Images of devices with increasing channel lengths (width was fixed at 45 μm). (e) Delivery efficiency and (f) viability 24 hours after cell stretching.

**Figure S6.**
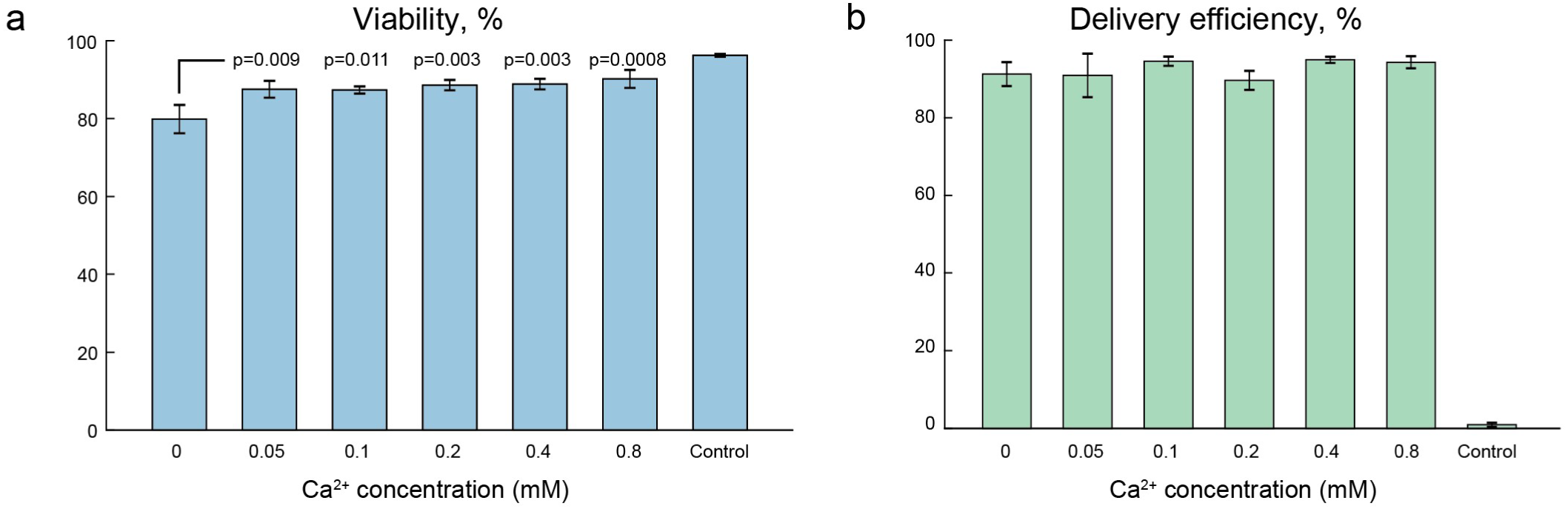
Addition of calcium ion improves same-day viability. Jurkat cells were suspended in transfection buffer of PBS with 2 mg/mL HA, 0.2 mg/mL FITC-Dextran 70 kDa, and varying amounts of Ca2+, and processed at 3 bar in contraction-only devices (e.g. Figure S1) 45 μm wide and 100 μm long, N=3 replicates for each condition. Viability in conditions containing calcium were each compared to the no-calcium condition by Tukey’s honest significance test, yielding indicated p-values.

**Figure S7.**
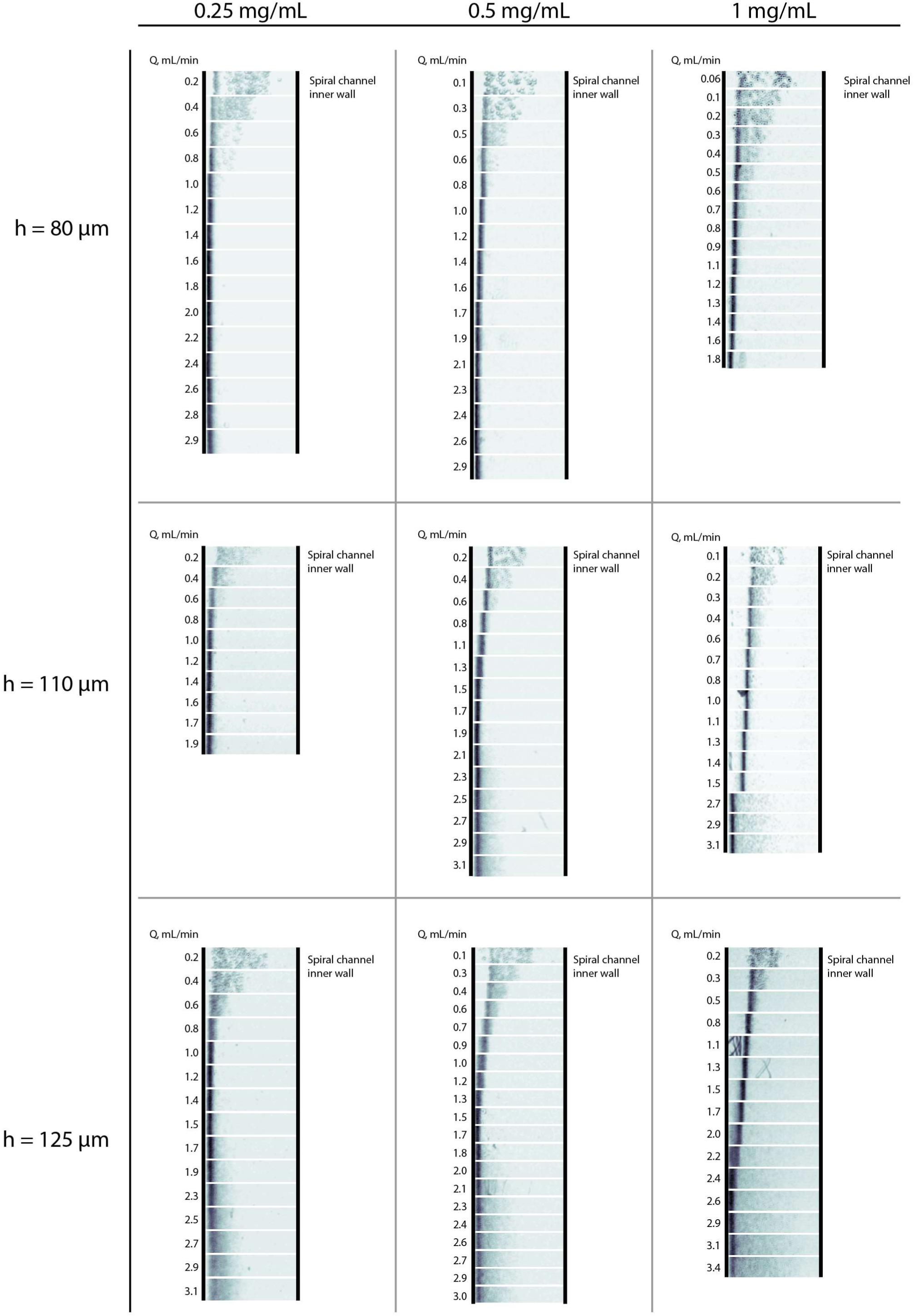
Composite timelapse images of cells in 500 µm wide spiral focusing channels, evaluated over a range of flow rates, for each combination of channel height and hyaluronic acid concentration.

**Figure S8.**
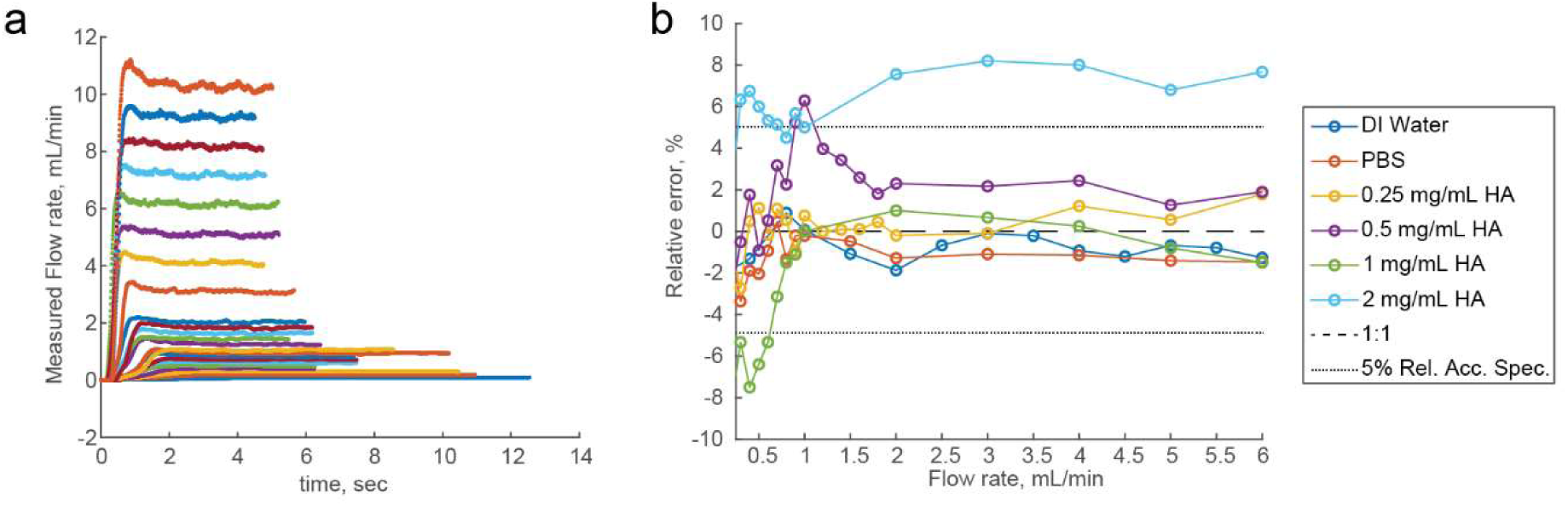
Validation of the heat pulse flow meter (Sensirion SLF35-1300F) for viscoelastic solutions. Solutions of 1.6 MDa HA in PBS at concentrations of 0 (none), 0.25 mg/mL, 0.5 mg/mL, 1 mg/mL, and 2 mg/mL. Solutions were driven using a precision syringe pump (Harvard Apparatus) through the flow meter at a flow rates from 0.1 to 10 mL/min. (a) Representative traces of measured flow rates vs. time revealed the pumping equilibration time of 2-5 seconds. (b) Tukey mean difference plot of the relative error between measured and prescribed flow rates. Dotted lines indicate the 5% relative accuracy specification of the flow meter. Measured flow rates for 2 mg/mL HA are systematically overestimated due to shear thinning.

**Figure S9.**
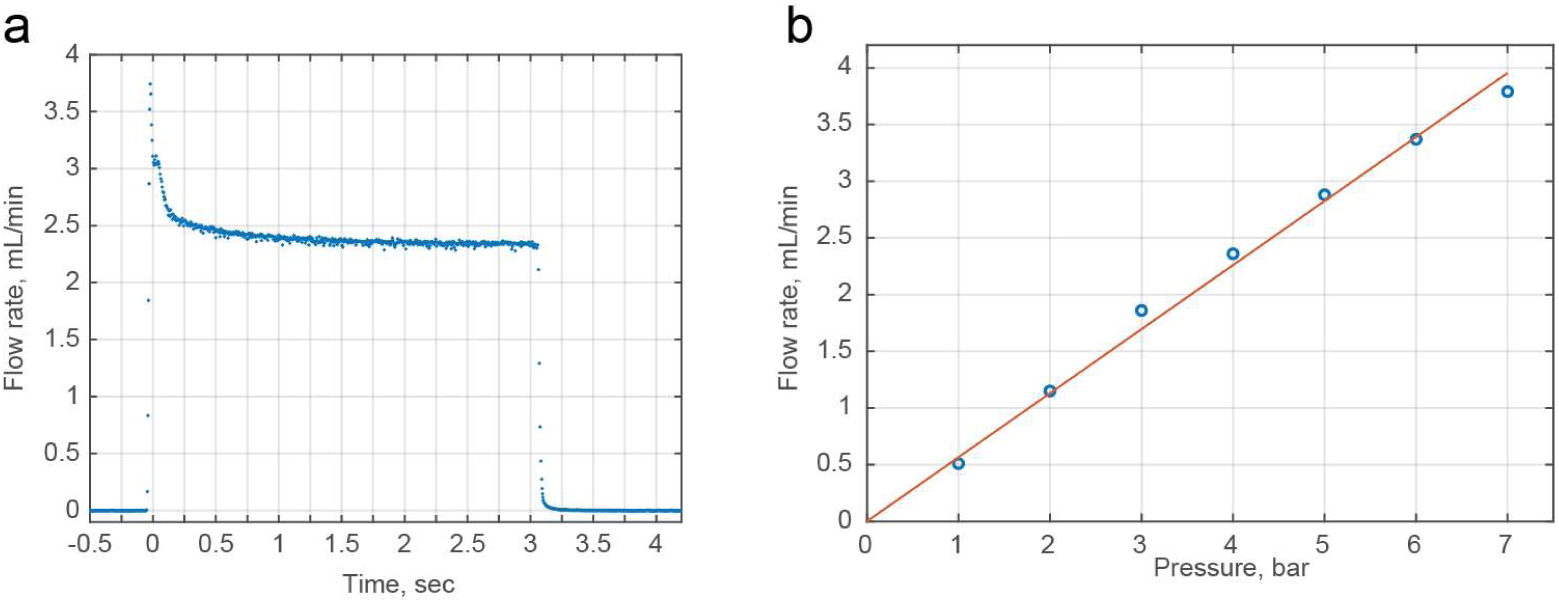
Flow rate vs. driving pressure for a 1 mg/mL HA solution in the chip with cell focusing channels. (a) Representative time trace of measured flow rate upon opening the pneumatic valve (4 bar). (b) Measured flow rates at pressures from 1 bar to 7 bar with linear regression, Q [mL/min] = 0.565 * P [bar].

**Figure S10.**
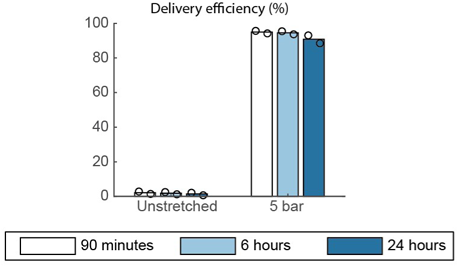
FITC-Dextran retention. Percent of Jurkat cells staining positive for 70 kDa FITC-Dextran after 90 minutes, 6 hours, and 24 hours post-transfection, as compared to unstretched controls.

**Figure S11.**
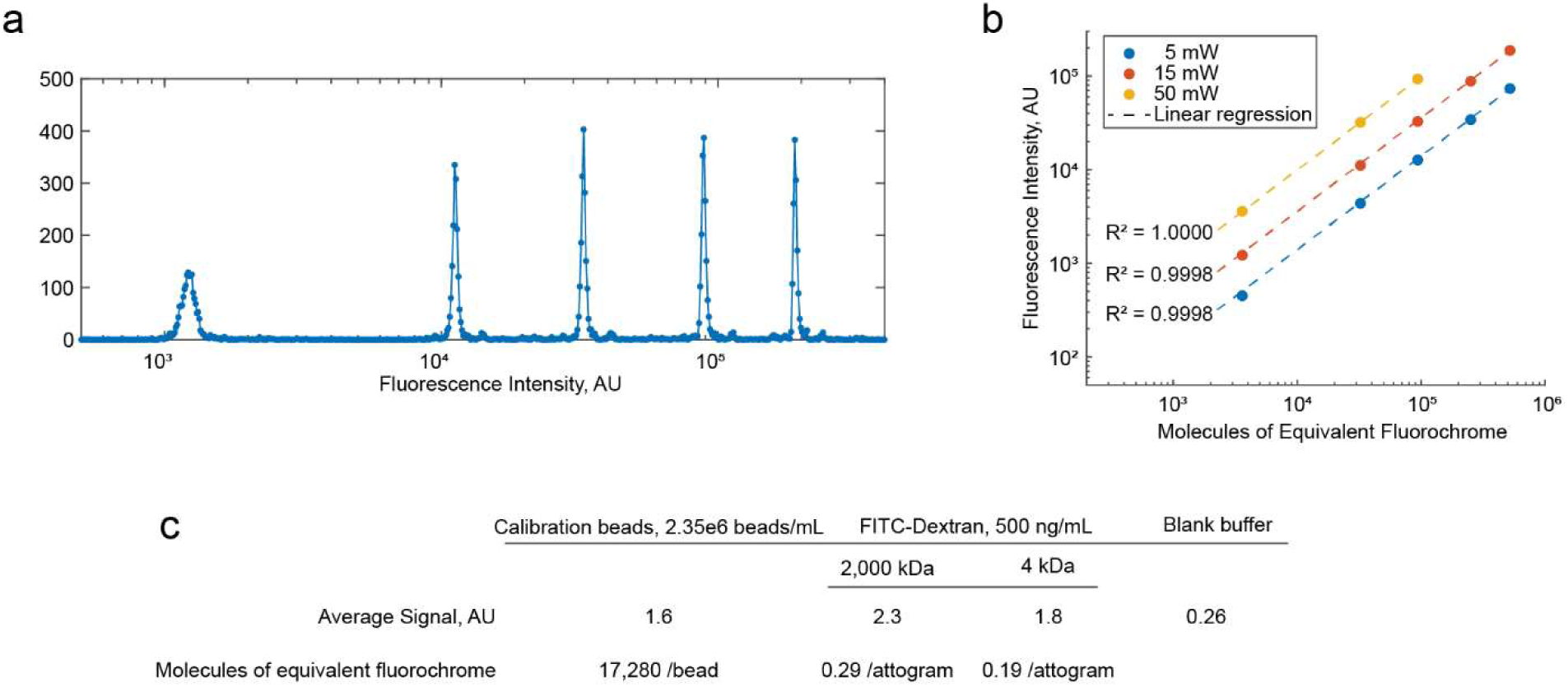
Flow cytometer calibration for quantitative measurements of dextran delivery. (a) Fluorescence intensity histogram of the calibration bead mixture in the FITC channel of the flow cytometer with an illumination intensity of 15 mW. (b) Calibration bead peak intensities plotted against MEF—molecules of equivalent fluorochrome—for FITC of each bead population, as provided by the manufacturer, at relevant cytometer laser powers. Linear regressions and goodness of fit were calculated on the raw (i.e., not log-transformed) data.

**Figure S12.**
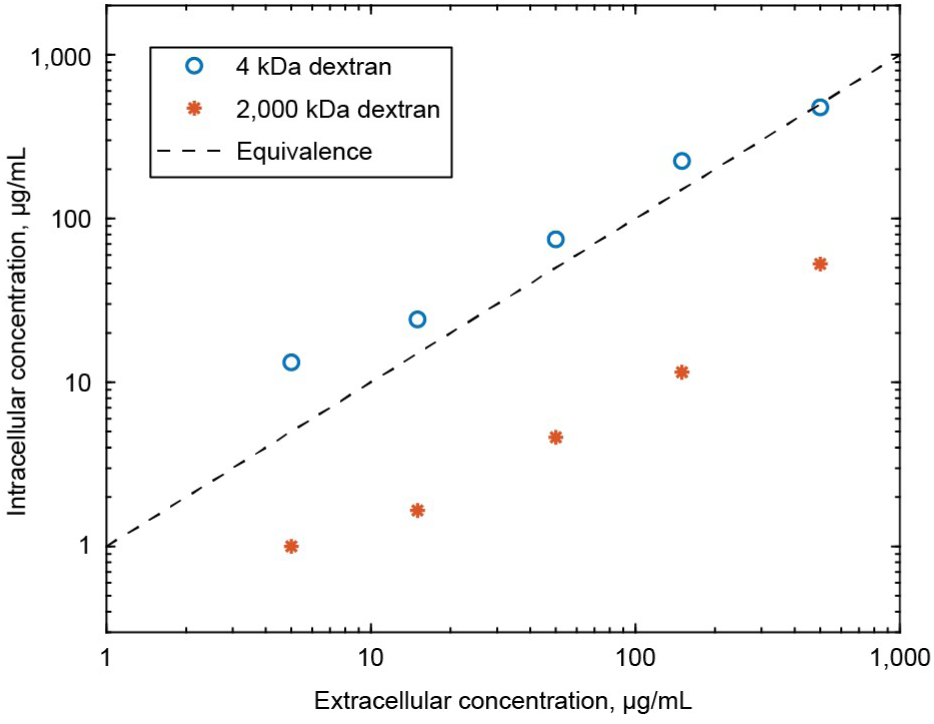
Estimated intracellular concentrations of small and large dextrans. Based on a nominal cell diameter of 12 μm (Figure S2) and assuming a spherical cell volume, the estimated intracellular concentrations for 4 kDa dextran were within a factor of 2 of the extracellular concentrations in the transfection solution, and about 10-fold less for 2,000 kDa dextran.

**Figure S13.**
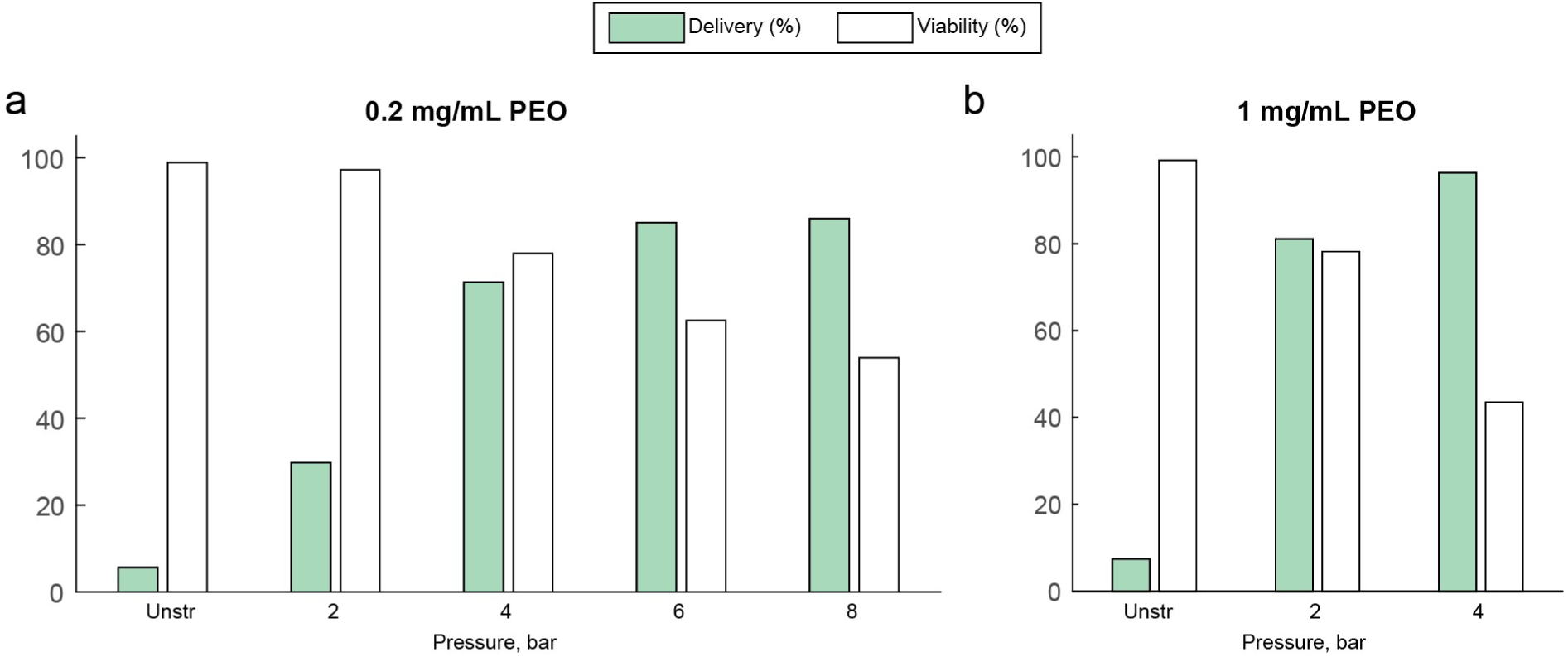
Generalization of viscoelastic mechanoporation beyond hyaluronic acid. Transfection of 70 kDa FITC-dextran to Jurkat cells in a transfection solution containing (a) 0.2 mg/mL or (b) 1 mg/mL of 2 MDa polyethylene oxide PEO) instead of HA. N=1, delivery & viability evaluated on the same day as transfection.

**Figure S14.**
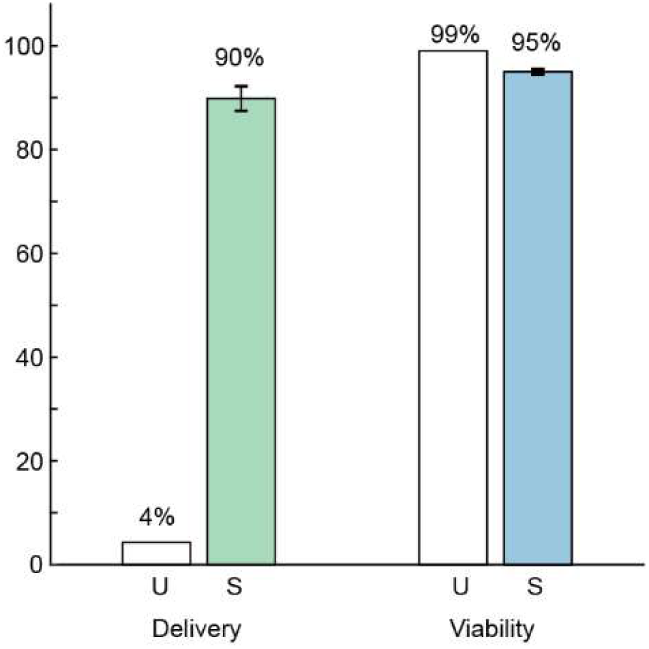
Efficient intracellular delivery of a protein to Jurkat cells. FITC-albumin at a concentration of 0.2 mg/mL was delivered by viscoelastic mechanoporation. Delivery efficiency and viability were evaluated 90 minutes after transfection. U, unstretched cells co-incubated with the FITC-albumin but not processed through the chip (n=1); S, ‘stretched’ cells processed through the chip (n=3).

**Figure S15.**
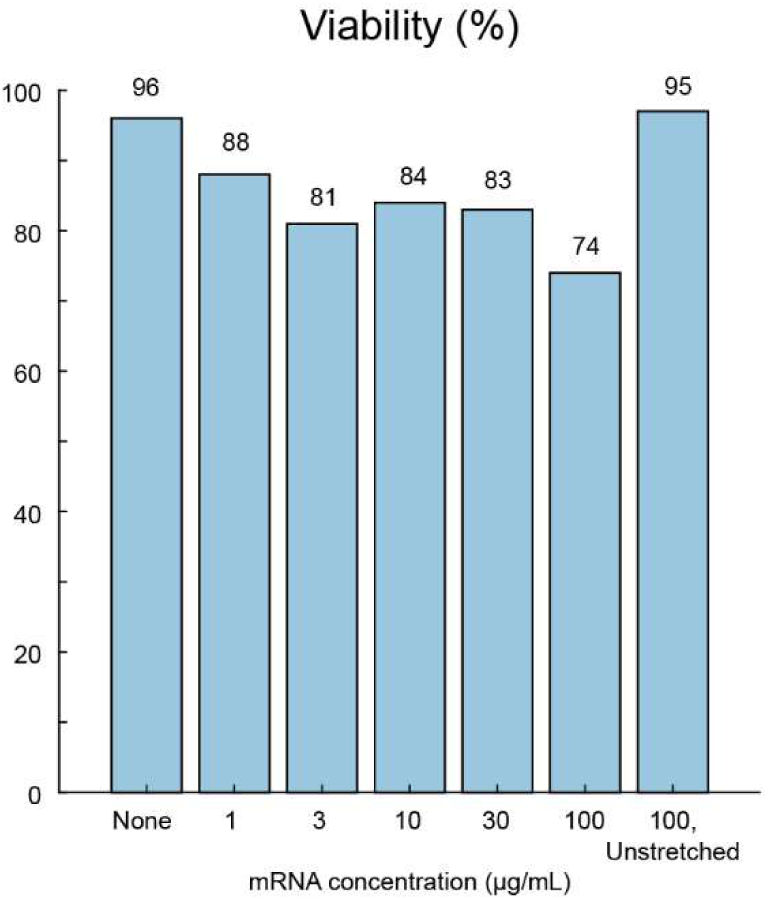
Viability of Jurkat cells following mRNA delivery. Viability was assessed by propidium iodide exclusion 24 hours after transfection.

**Figure S16.**
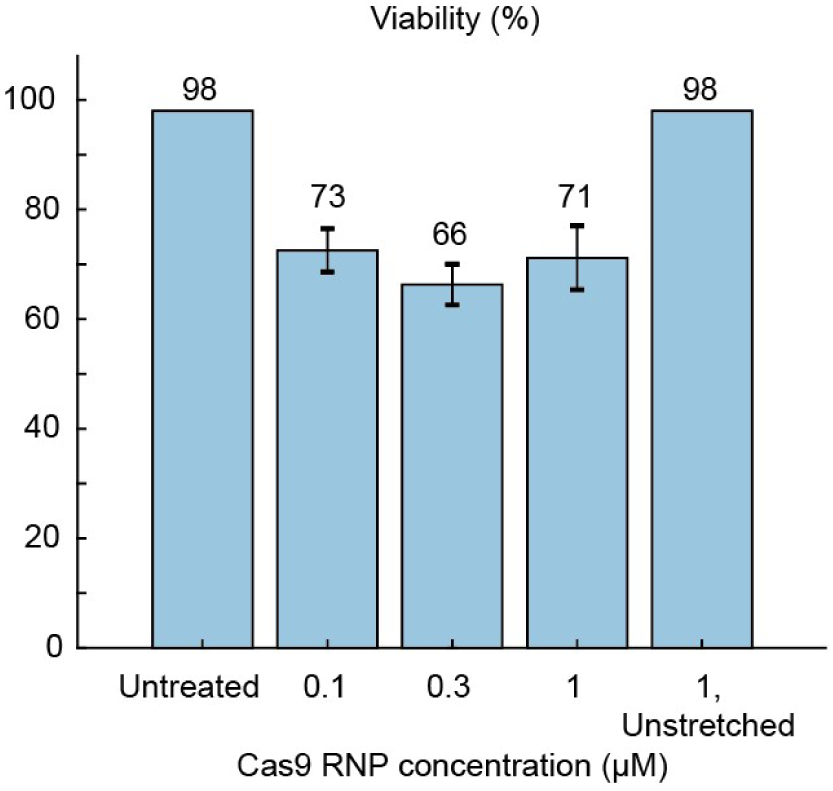
Viability of Jurkat cells 2 days after RNP delivery. Viability was assessed by propidium iodide exclusion.

**Figure S17.**
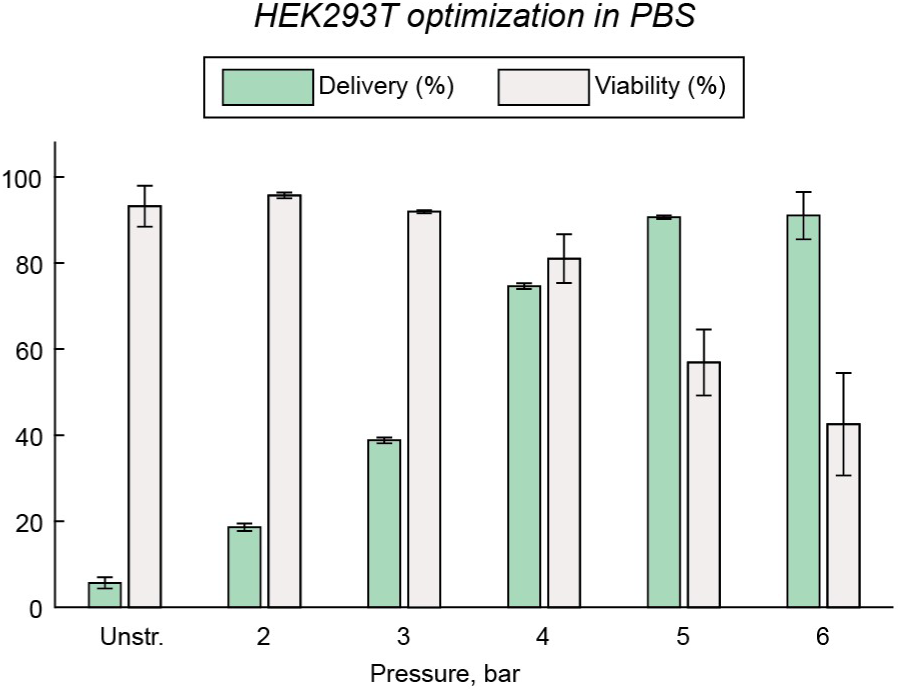
Optimization of HEK293T cell transfection with 70 kDa FITC-dextran. Dextran delivery and cell viability about 90 minutes after processing through the chip with PBS-based transfection buffer. N=3 replicates per condition.

**Figure S18.**
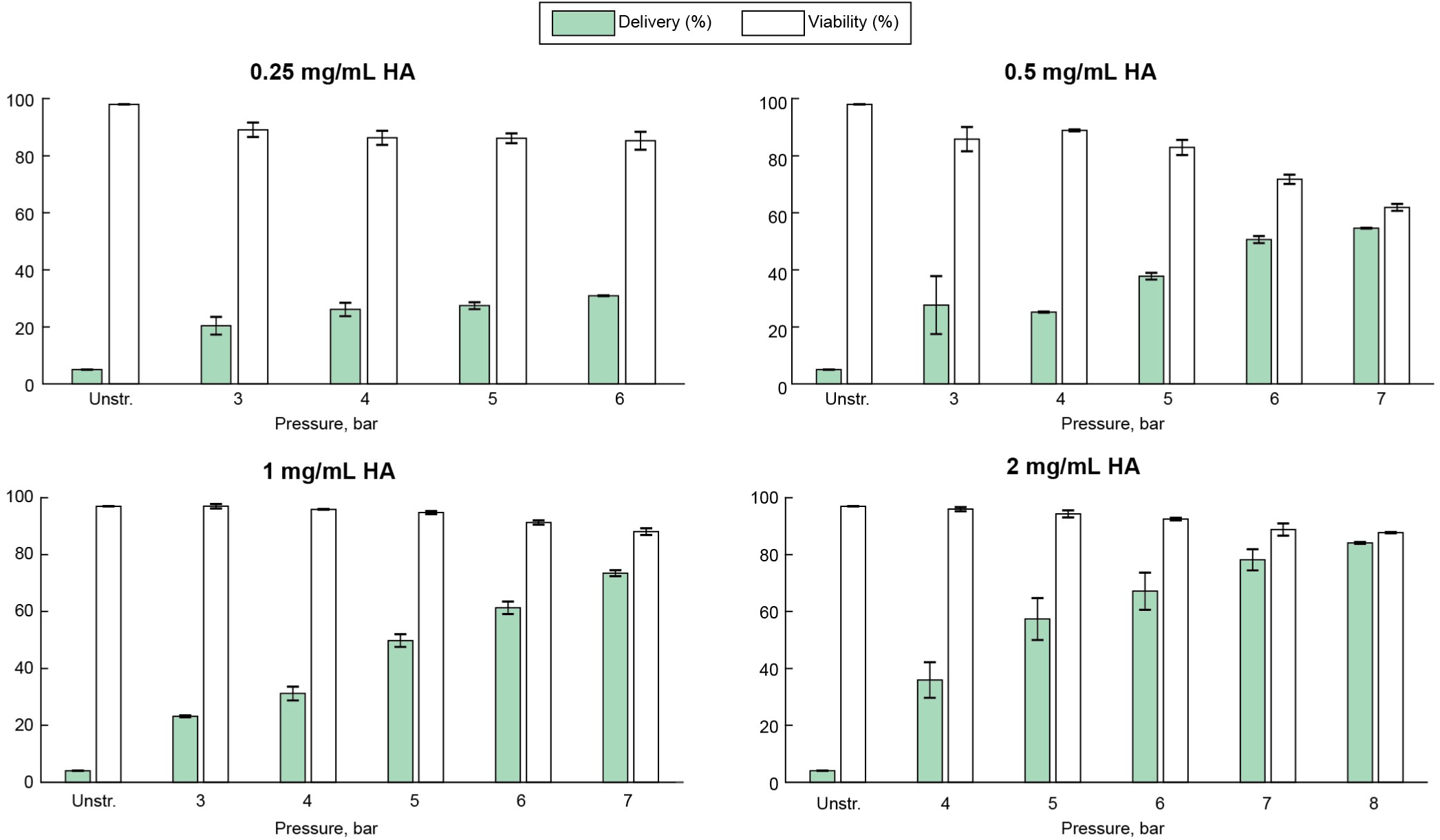
Optimization of primary T cell transfection with 70 kDa FITC-dextran. Delivery efficiency and viability were evaluated the same day following transfection with a range of driving pressures for delivery solutions containing 0.25 mg/mL, 0.5 mg/mL, 1 mg/mL or 2 mg/mL HA. N=2 replicates per condition.

**Figure S19.**
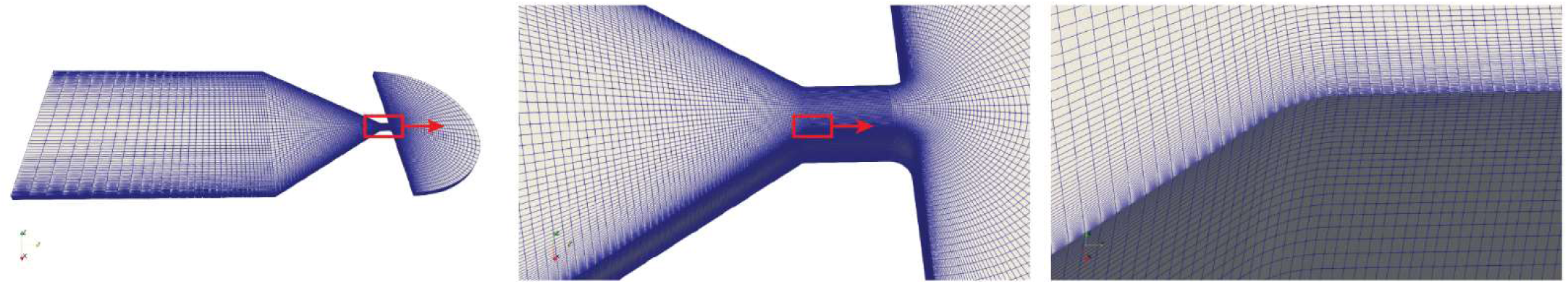
Detailed views of the mesh used for inertio-elastic flow simulations.

**Figure S20.**
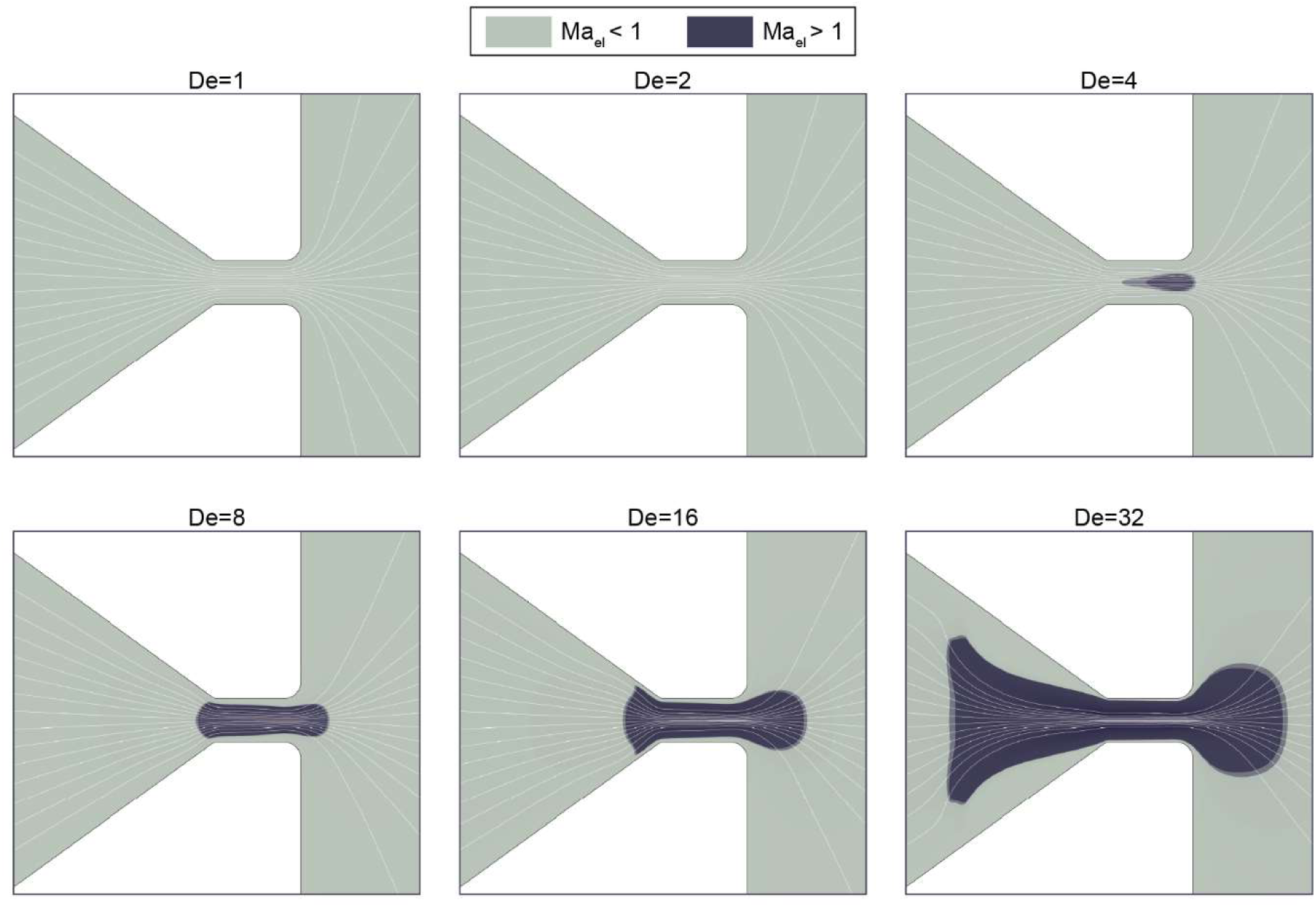
Deviations from the laminar flow profile and upstream flow separation are associated with the local viscoelastic Mach number exceeding one in computational simulations. Regions where the local viscoelastic Mach number is either smaller (green) or larger (blue) than one are visualized for increasing Deborah numbers (described in text). Streamlines show deviations from laminar flow starting at De=4 and upstream separation at De=32, coinciding with local transitions to 𝑀𝑎_el_> 1.

**Table S1.**
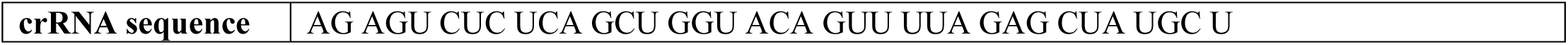
crRNA sequence selected for TCR knockout study. crRNA was synthesized by Integrated DNA technologies and included both 5’ and 3’ DT Alt-R® end-blocking modifications.

## Viscoelastic effects within a contracting channel

The mechanoporation component of the device was a microfluidic contraction, which generated a uniaxial extensional flow along the channel centerline. In an idealized uniaxial extensional flow, the fluid deformation can be considered shear-free, such that the deformation rate 𝛾̇ is:

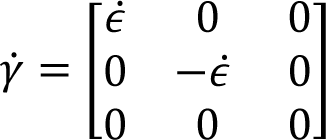

This results in fluid stresses that are related to the strain rate by a quantity called the extensional viscosity 𝜂_e_ such that 𝜏_xx_ = 𝜂_e_𝜖̇. For (generalized) Newtonian fluids, the extensional viscosity is geometrically fixed to be three times the shear viscosity, such that the ratio of viscosities or Trouton Ratio 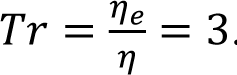. However, the Tr of viscoelastic flows can greatly exceed this, e.g. 𝑇𝑟 = 100 or even higher. At the molecular level, the increased viscosity of the HA solution is caused by the change in entropy associated with increasing the extension and alignment of the glycosaminoglycan chains. Because of this, the extensional viscosity is not a material constant but will typically increase with increasing fluid extension and chain alignment, a phenomenon known as strain hardening. In this work, the contraction ratio (i.e., the ratio of the upstream channel cross sectional area to the area at the narrowest point) was made as large as practical to maximize strain hardening.

To contextualize the experimental and computational studies of the upstream vortices, we briefly consider the viscoelastic Mach number, 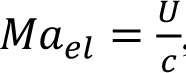, a dimensionless group comparing the flow velocity 𝑈 to the elastic wave speed 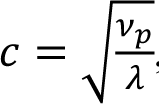, where is the polymer contribution to the zero-shear kinematic viscosity.^1^ When the polymer contribution to viscosity is much greater than that of the solvent (as is the case here), 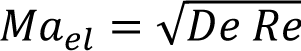, where 𝑅𝑒 is the well-known Reynolds number *Re* = 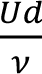. The transition to 𝑀𝑎 > 1 has been previously observed to be important for the onset of upstream flow instabilities for accelerating viscoelastic solutions.^1, 2^ Indeed, the onset and extent of flow separation in the computational results was observed to coincide with regions where the local 𝑀𝑎_el_ became greater than one in upstream contraction (**Figure S20**).

## References

1. Dobrowsky, T., Gianni, D., Pieracci, J. & Suh, J. AAV manufacturing for clinical use: Insights on current challenges from the upstream process perspective. Curr. Opin. Biomed. Eng. 20, 100353 (2021).

2. Dwarshuis, N. J., Parratt, K., Santiago-Miranda, A. & Roy, K. Cells as advanced therapeutics: State-of-the-art, challenges, and opportunities in large scale biomanufacturing of high-quality cells for adoptive immunotherapies. Adv. Drug Deliv. Rev. 114, 222–239 (2017).

3. Levine, B. L., Miskin, J., Wonnacott, K. & Keir, C. Global Manufacturing of CAR T Cell Therapy. Mol. Ther. - Methods Clin. Dev. 4, 92–101 (2017).

4. Lee, C. S. et al. Adenovirus-mediated gene delivery: Potential applications for gene and cell-based therapies in the new era of personalized medicine. Genes Dis. 4, 43–63 (2017).

5. Milone, M. C. & O’Doherty, U. Clinical use of lentiviral vectors. Leukemia 32, 1529–1541 (2018).

6. Marcucci, K. T. et al. Retroviral and Lentiviral Safety Analysis of Gene-Modified T Cell Products and Infused HIV and Oncology Patients. Mol. Ther. 26, 269–279 (2018).

7. Lächelt, U. & Wagner, E. Nucleic Acid Therapeutics Using Polyplexes: A Journey of 50 Years (and Beyond). Chem. Rev. 115, 11043–11078 (2015).

8. van der Loo, J. C. M. & Wright, J. F. Progress and challenges in viral vector manufacturing. Hum. Mol. Genet. 25, R42–R52 (2016).

9. Stewart, M. P., Langer, R. & Jensen, K. F. Intracellular Delivery by Membrane Disruption: Mechanisms, Strategies, and Concepts. Chem. Rev. 118, 7409–7531 (2018).

10. Weaver, J. C. Electroporation of biological membranes from multicellular to nano scales. IEEE Trans. Dielectr. Electr. Insul. 10, 754–768 (2003).

11. Gowrishankar, T. R., Stern, J. V. & Weaver, J. C. Electroporation dynamics for different pore lifetimes based on the standard model. ArXiv171003650 Phys. (2017).

12. Meaking, W. S., Edgerton, J., Wharton, C. W. & Meldrum, R. A. Electroporation-induced damage in mammalian cell DNA. Biochim. Biophys. Acta BBA - Gene Struct. Expr. 1264, 357–362 (1995).

13. Maccarrone, M., Rosato, N. & Agro, A. F. Electroporation Enhances Cell Membrane Peroxidation and Luminescence. Biochem. Biophys. Res. Commun. 206, 238–245 (1995).

14. Hui, S. W. & Li, L. H. In Vitro and Ex Vivo Gene Delivery to Cells by Electroporation. in Electrochemotherapy, Electrogenetherapy, and Transdermal Drug Delivery: Electrically Mediated Delivery of Molecules to Cells (eds. Jaroszeski, M. J., Heller, R. & Gilbert, R.) 157– 171 (Humana Press, 2000). doi:10.1385/1-59259-080-2:157.

15. Gabriel, B. & Teissié, J. Generation of reactive-oxygen species induced by electropermeabilization of Chinese hamster ovary cells and their consequence on cell viability. Eur. J. Biochem. 223, 25–33 (1994).

16. Lissandrello, C. A. et al. High-throughput continuous-flow microfluidic electroporation of mRNA into primary human T cells for applications in cellular therapy manufacturing. Sci. Rep. 10, 18045 (2020).

17. Sido, J. M. et al. Electro-mechanical transfection for non-viral primary immune cell engineering. 2021.10.26.465897 https://www.biorxiv.org/content/10.1101/2021.10.26.465897v1(2021) doi:10.1101/2021.10.26.465897.

18. Meacham, J. M., Durvasula, K., Degertekin, F. L. & Fedorov, A. G. Physical Methods for Intracellular Delivery: Practical Aspects from Laboratory Use to Industrial-Scale Processing. J. Lab. Autom. 19, 1–18 (2014).

19. Evans, E., Heinrich, V., Ludwig, F. & Rawicz, W. Dynamic Tension Spectroscopy and Strength of Biomembranes. Biophys. J. 85, 2342–2350 (2003).

20. Sharei, A. et al. A vector-free microfluidic platform for intracellular delivery. Proc. Natl. Acad. Sci. 201218705 (2013) doi:10.1073/pnas.1218705110.

21. Liu, A. et al. Microfluidic generation of transient cell volume exchange for convectively driven intracellular delivery of large macromolecules. Mater. Today 21, 703–712 (2018).

22. Hur, J. et al. Microfluidic Cell Stretching for Highly Effective Gene Delivery into Hard-to-Transfect Primary Cells. ACS Nano (2020) doi:10.1021/acsnano.0c05169.

23. Dixit, H. G. et al. Massively-Parallelized, Deterministic Mechanoporation for Intracellular Delivery. Nano Lett. 20, 860–867 (2020).

24. Nejadnik, H. et al. Instant labeling of therapeutic cells for multimodality imaging. Theranostics 10, 6024–6034 (2020).

25. Liu, A. et al. Cell Mechanical and Physiological Behavior in the Regime of Rapid Mechanical Compressions that Lead to Cell Volume Change. Small 16, 1903857 (2020).

26. Kwon, C. & Chung, A. J. Highly efficient mRNA delivery with nonlinear microfluidic cell stretching for cellular engineering. Lab. Chip (2023) doi:10.1039/D2LC01115H.

27. Uvizl, A. et al. Efficient and gentle delivery of molecules into cells with different elasticity via Progressive Mechanoporation. Lab. Chip 21, 2437–2452 (2021).

28. Adamo, A. & Jensen, K. F. Microfluidic based single cell microinjection. Lab. Chip 8, 1258– 1261 (2008).

29. Meacham, J. M., Durvasula, K., Degertekin, F. L. & Fedorov, A. G. Enhanced intracellular delivery via coordinated acoustically driven shear mechanoporation and electrophoretic insertion. Sci. Rep. 8, 3727 (2018).

30. Belling, J. N. et al. Acoustofluidic sonoporation for gene delivery to human hematopoietic stem and progenitor cells. Proc. Natl. Acad. Sci. 117, 10976–10982 (2020).

31. Hallow, D. M. et al. Shear-induced intracellular loading of cells with molecules by controlled microfluidics. Biotechnol. Bioeng. 99, 846–854 (2008).

32. Deng, Y. et al. Intracellular Delivery of Nanomaterials via an Inertial Microfluidic Cell Hydroporator. Nano Lett. 18, 2705–2710 (2018).

33. Joo, B., Hur, J., Kim, G.-B., Yun, S. G. & Chung, A. J. Highly Efficient Transfection of Human Primary T Lymphocytes Using Droplet-Enabled Mechanoporation. ACS Nano (2021) doi:10.1021/acsnano.0c10473.

34. Kizer, M. E. et al. Hydroporator: a hydrodynamic cell membrane perforator for high-throughput vector-free nanomaterial intracellular delivery and DNA origami biostability evaluation. Lab. Chip (2019) doi:10.1039/C9LC00041K.

35. Jarrell, J. A. et al. Intracellular delivery of mRNA to human primary T cells with microfluidic vortex shedding. Sci. Rep. 9, 3214 (2019).

36. Jarrell, J. A. et al. Numerical optimization of microfluidic vortex shedding for genome editing T cells with Cas9. Sci. Rep. 11, 11818 (2021).

37. Lokhandwalla, M. & Sturtevant, B. Mechanical haemolysis in shock wave lithotripsy (SWL): I. Analysis of cell deformation due to SWL flow-fields. Phys. Med. Biol. 46, 413–437 (2001).

38. Boucher, P.-A., Joós, B., Zuckermann, M. J. & Fournier, L. Pore Formation in a Lipid Bilayer under a Tension Ramp: Modeling the Distribution of Rupture Tensions. Biophys. J. 92, 4344– 4355 (2007).

39. Razizadeh, M., Nikfar, M., Paul, R. & Liu, Y. Coarse-Grained Modeling of Pore Dynamics on the Red Blood Cell Membrane under Large Deformations. Biophys. J. 119, 471–482 (2020).

40. Moe, A. M., Golding, A. E. & Bement, W. M. Cell healing: Calcium, repair and regeneration. Semin. Cell Dev. Biol. 45, 18–23 (2015).

41. Davenport, N. R. & Bement, W. M. Cell repair: Revisiting the patch hypothesis. Commun. Integr. Biol. 9, e1253643 (2016).

42. Jimenez, A. J. et al. ESCRT Machinery Is Required for Plasma Membrane Repair. Science 343, 1247136 (2014).

43. Xiang, N. et al. Fundamentals of elasto-inertial particle focusing in curved microfluidic channels. Lab. Chip 16, 2626–2635 (2016).

44. Lee, H. S. & Muller, S. J. A differential pressure extensional rheometer on a chip with fully developed elongational flow. J. Rheol. 61, 1049–1059 (2017).

45. Roth, T. L. et al. Reprogramming human T cell function and specificity with non-viral genome targeting. Nature 559, 405 (2018).

46. Hendel, A. et al. Chemically modified guide RNAs enhance CRISPR-Cas genome editing in human primary cells. Nat. Biotechnol. 33, 985–989 (2015).

47. Kang, G. et al. Intracellular Nanomaterial Delivery via Spiral Hydroporation. ACS Nano (2020) doi:10.1021/acsnano.9b07930.

48. Lee, S. S., Yim, Y., Ahn, K. H. & Lee, S. J. Extensional flow-based assessment of red blood cell deformability using hyperbolic converging microchannel. Biomed. Microdevices 11, 1021 (2009).

49. Alves, M. A., Oliveira, P. J. & Pinho, F. T. Numerical Methods for Viscoelastic Fluid Flows. Annu. Rev. Fluid Mech. 53, 509–541 (2021).

50. Rein, C., Toner, M. & Sevenler, D. Rapid prototyping for high-pressure microfluidics. Sci. Rep. 13, 1–9 (2023).

51. Ronneberger, O., Fischer, P. & Brox, T. U-Net: Convolutional Networks for Biomedical Image Segmentation. in Medical Image Computing and Computer-Assisted Intervention – MICCAI 2015 (eds. Navab, N., Hornegger, J., Wells, W. M. & Frangi, A. F.) 234–241 (Springer International Publishing, 2015). doi:10.1007/978-3-319-24574-4_28.

52. Pimenta, F. & Alves, M. A. Stabilization of an open-source finite-volume solver for viscoelastic fluid flows. J. Non-Newton. Fluid Mech. 239, 85–104 (2017).

53. Krause, W. E., Bellomo, E. G. & Colby, R. H. Rheology of Sodium Hyaluronate under Physiological Conditions. Biomacromolecules 2, 65–69 (2001).

54. Bingöl, Ö., Lohmann, D., Püschel, K. & Kulicke, W.-M. Characterization and comparison of shear and extensional flow of sodium hyaluronate and human synovial fluid. Biorheology 47, 205–24 (2010).

55. Haward, S. J., Jaishankar, A., Oliveira, M. S. N., Alves, M. A. & McKinley, G. H. Extensional flow of hyaluronic acid solutions in an mnoptimized microfluidic cross-slot device. Biomicrofluidics 7, 044108 (2013).

56. Herrchen, M. & Öttinger, H. C. A detailed comparison of various FENE dumbbell models. J. Non-Newton. Fluid Mech. 68, 17–42 (1997).

## References

1. Rodd, L. E., Cooper-White, J. J., Boger, D. V. & McKinley, G. H. Role of the elasticity number in the entry flow of dilute polymer solutions in micro-fabricated contraction geometries. Journal of Non-Newtonian Fluid Mechanics 143, 170–191 (2007).

2. Shi, X. & Christopher, G. F. Growth of viscoelastic instabilities around linear cylinder arrays. Physics of Fluids 28, 124102 (2016).

